# Criterion-selective neurons in the human medial frontal cortex track decision thresholds during memory-based decision making

**DOI:** 10.64898/2026.08.19.745781

**Authors:** Evan Layher, Ivan Skelin, Chrystal M. Reed, Jeffrey M. Chung, Lisa M. Bateman, Taufik A. Valiante, Adam N. Mamelak, Michael B. Miller, Ueli Rutishauser

**Author notes:** These authors contributed equally to this work (Michael B. Miller, Ueli Rutishauser).

## Abstract

The decision criterion is foundational to theories of decision-making, yet little is known about its neural underpinnings. In memory-based decision making, the criterion sets the minimal memory strength for something to be familiar, but whether or how it is distinctly represented from memory strength is unknown. We recorded single neurons in the medial frontal cortex (MFC) and medial temporal lobe (MTL), both implicated in memory-based decisions, while participants made decisions under different decision criteria. We identified criterion-selective (CS) neurons in the MFC that tracked the criterion regardless of memory strength, and memory-selective (MS) neurons in both regions that tracked memory strength regardless of the criterion. CS neurons signaled the criterion before MS neurons signaled memory strength, and a race model incorporating both neuron types outperformed one using MS neurons alone. These findings reveal two independent cellular substrates, one for the decision criterion and one for memory strength, whose joint activity underlies memory-based decisions.

## Introduction

Every day we make decisions based on our memories, yet our memories are often uncertain. Consider the common experience of encountering a potentially familiar face: do you say “hello” and risk embarrassing yourself in front of a stranger, or stay silent and risk offending a known acquaintance? This decision hinges on two factors: the strength of the memory evoked by that face, and a threshold determining the minimum amount of memory strength we consider sufficient to decide that the face is familiar—the decision criterion. The concept of the decision criterion is foundational to theories of decision-making (*1–4*). In particular, it is a core component of signal detection theory (SDT), a framework widely used to model decision-making (*5*). Importantly, the decision criterion is not fixed. Rather, the criterion needs to be set flexibly in response to situational demands, a capacity known as criterion shifting that is essential for making decisions (*6*). For instance, a presumed co-worker encountered at work warrants recognition even under weak memory evidence (a lax criterion), given the high likelihood of the encounter. However, the same face spotted unexpectedly while on vacation abroad demands much stronger evidence before recognition (a strict criterion), given the improbability of such a meeting. Despite its importance in decision-making generally, and in memory-based decisions specifically, little is known about the neural mechanisms by which a decision criterion is represented distinctly from the underlying signal. Single-unit investigations of the criterion during perceptual decision-making in macaques have identified neurons that *integrate* noisy sensory evidence with a decision criterion to guide choices, in the lateral intraparietal area (*7–11*) and frontal eye field (*12*). While these findings show that criterion shifts can modulate neuronal activity during sensory processing, they leave open a more fundamental question: how is the decision criterion itself represented, independently of the signal it is applied to? Here, we identify a single-neuron correlate in the human medial frontal cortex (MFC) that independently tracks the decision criterion during memory-based decision-making.

The neural basis of memory has long been associated with the medial temporal lobe (MTL), a view cemented by the 1953 bilateral MTL resection in patient H.M., which left him unable to form new episodic memories (*13*). Notably, H.M. retained many memories from before the surgery, indicating that certain memory processes extend beyond the MTL. With the advent of functional magnetic resonance imaging (fMRI) in the 1990s, it became apparent that regions throughout the cerebral cortex also contribute to memory (*14*, *15*). Among these, the MFC emerged as particularly important in memory-based decision-making (*14*, *16*), in addition to its well-established roles in cognitive control (*17*), performance monitoring (*18*), and flexible decision-making (*19*). Since the MFC supports multiple cognitive processes, it has remained difficult to disentangle its role in memory from that in decisional processes (*20*). For example, while fMRI studies of recognition memory consistently show differential MFC activity between previously studied (old) and novel (new) choices (old > new choice contrast), it remains unclear whether these differences reflect memory (*21–26*), decisional processes (*27–33*), or a mixture thereof. Since fMRI results are mixed (*21–33*) and individual MFC neurons encode different cognitive processes without apparent anatomical clustering (*16*, *18*), we posit that at the single neuron level, memory and decisional processes can be differentiated in the MFC. Addressing this major gap requires an experiment in which both the decision criterion and memory strength are independently manipulated to separate the two processes, which we do here.

Given the ample evidence implicating the MFC in memory-based decision-making (*14*, *16*, *20–33*), we predicted that the decision criterion would be represented in a subset of MFC neurons. Motivated by SDT (Fig. 1A), we hypothesized that neurons encoding the criterion should modulate their firing rates when the criterion shifts, while remaining insensitive to memory strength. Our first major contribution is to identify and characterize such neurons in the MFC.

**Fig. 1.**
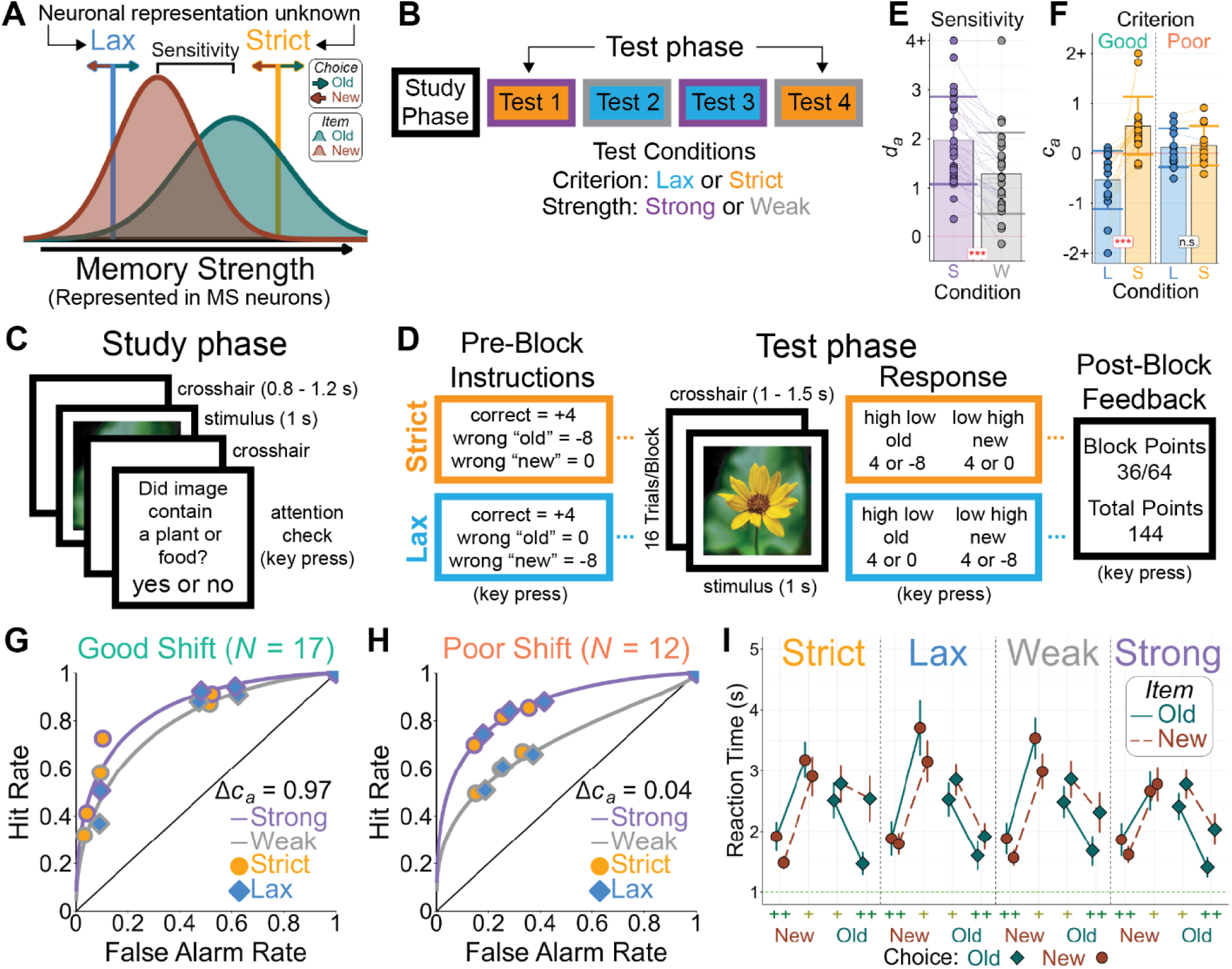
Task and behavior. (**A**) Signal Detection Theory (SDT) model of recognition memory. Old (green) and new (red) item distributions are represented as overlapping Gaussians along a continuous memory strength axis, whose separation reflects sensitivity. Lax (blue) and strict (orange) decision criteria are shown, indicating the minimum memory strength required to classify an item as old. Memory-selective (MS) neurons are known to encode memory strength, but single neuron correlates of the decision criterion remain unknown. (**B**) Task structure. Participants completed a study phase in which novel images were presented once or twice to induce weak or strong memory strength, followed by four test blocks crossing memory strength (weak [gray], strong [purple]) and criterion (strict [orange], lax [blue]) conditions. This study/test cycle repeated four times, with the order of test blocks randomized. (**C**) Study phase. Participants studied 16 images once and 16 twice, with attention checks following 8 randomly selected trials. (**D**) Test phase. Participants were instructed on the point values assigned to each choice type, then judged images as old or new with confidence ratings to maximize their total points. Block-level and cumulative point totals were displayed after each block. (**E**) Mean sensitivity (d_a_) ±SD showing significantly greater sensitivity in the strong (purple, S) than weak (gray, W) condition. (**F**) Mean decision criteria (c_a_) ±SD for good (left) and poor (right) shifters across lax (blue, L) and strict (orange, S) criterion conditions. Good shifters showed a significant difference in criteria between conditions; poor shifters did not. (**G** and **H**) ROC curves for good (**G**) and poor (**H**) shifters, showing aggregated performance across conditions. (**I**) Mean reaction times (±SEM) by choice type and condition. Responses were faster for high confidence (++, green) than low confidence (+, yellow) choices, and for correct than incorrect responses, across all conditions.

Single-unit recordings during recognition memory have revealed a cellular substrate for memory strength in the form of memory-selective (MS) neurons, which fire differentially to correctly identified old (hits) versus new (correct rejections) items (H > CR). MS neurons have been identified in both the MFC (*16*) and MTL (*16*, *34*). In the MTL, their firing rates scale with decision confidence (*34*), consistent with the notion that these cells track memory strength, as confidence is thought to reflect the strength of a memory trace (*35*). A core assumption of SDT is that memory strength and criterion placement are independent processes (*6*), and thus should be supported by distinct neuronal populations. Our second major contribution is a direct test of this assumption at the cellular level. We do so by examining MS neuron activity under different criterion settings and contrasting it with activity of neurons that signal the criterion.

Finally, examining decision criteria is complicated by substantial individual differences in the degree to which individuals shift their criteria, even when explicitly aware of the advantages. Some individuals shift criteria effectively, while others fail to shift entirely, despite clear instructions (*36*). These tendencies are consistent across criterion manipulation types (*37*, *38*), task contexts (*39*), decision domains (*40*, *41*), and time (*41*), yet are unrelated to personality traits, motivation, or other cognitive abilities (*37*, *39*, *41*). Importantly, individuals who fail to adequately shift their criteria show virtually no change in MFC activity across criterion conditions in fMRI (*27*), suggesting that neural effects related to the criterion are only detectable in those who shift sufficiently. We therefore predicted that single neuron correlates of the decision criterion would only be observable in individuals who adequately shift their criteria.

### Task and behavior

We developed a recognition memory paradigm in which the decision criterion and memory strength were each manipulated independently across two levels (Fig. 1B). The task consisted of two parts: a study phase, during which subjects viewed novel images to be remembered (Fig. 1C), followed by a recognition test phase (Fig. 1D). During the study phase, images were presented either once or twice to induce relatively weak or strong memories, respectively. The test phase consisted of four blocks, each covering a different combination of memory strength (weak vs. strong) and criterion (lax vs. strict). Before each test block, subjects saw instructions indicating whether to apply a lax or strict criterion for that block, followed by a series of 16 test images. For each image, participants decided whether it was “old” or “new” and rated their confidence as “high” or “low.” Confidence ratings were used to gauge memory strength, yielding four response categories ordered from weakest to strongest (high new < low new < low old < high old). The instructions preceding each test block informed subjects of the points earned or lost for each response type, as a way to manipulate the decision criterion. Correct responses earned 4 points, while false alarms and misses were designated as either critical (−8 points) or noncritical (no penalty) errors. False alarms served as critical errors in the strict condition, encouraging participants to favor “new” responses, while misses served as critical errors in the lax condition, encouraging them to favor “old” responses. After each test block, participants received feedback on points earned during that block and their running total.

Memory performance was assessed using the SDT measure *d_a_*, which indexes the separation in mean memory strength between “old” and “new” items (a quantity known as sensitivity) independently of the decision criterion (*6*). A *d_a_* value of 0 indicates chance performance, and increasingly positive values reflect increasingly higher sensitivity. As expected, sensitivity was greater in the strong than weak memory condition (strong: *M d_a_* = 1.97 ± 0.60; weak: *M d_a_* = 1.29 ± 0.36; *M* Δ*d_a_* = 0.68, 95% CI [0.22, 1.13]; Fig. 1E).

Criterion placement was assessed using the SDT measure *c_a_*, which indexes the decision criterion independently of sensitivity, where 0 represents a neutral criterion, negative values reflect a lax criterion, and positive values reflect a strict criterion (*6*). The difference in *c_a_* between the strict and lax conditions (Δ*c_a_*) served as a measure of criterion shifting, where zero indicates no shift occurred and positive values indicating the extent of the criterion shift. We divided subjects into two groups based on whether they shifted their criteria (good shifters: 17 sessions from 12 subjects; poor shifters: 12 sessions from 8 subjects), using a cutoff of Δ*c_a_* = 0.30 rather than zero to ensure that observed shifts reflected a meaningful change in decision strategy, consistent with prior research (*27*, *32*). This division was motivated by the expectation that neural signals related to criterion adjustment would only be detectable in subjects who genuinely changed their decision strategy (*27*). Good shifters adjusted their decision criteria between the strict (*M c_a_* = 0.55 ± 0.58) and lax (*M c_a_* = -0.53 ± 0.58) conditions (*M* Δ*c_a_* = 1.08, 95% CI [0.68, 1.49]), whereas poor shifters showed no significant change (strict: *M c_a_* = 0.15 ± 0.40; lax: *M c_a_* = 0.11 ± 0.39; *M* Δ*c_a_* = 0.04, 95% CI [-0.29, 0.38]; Fig. 1F). Overall, these results indicate that the behavioral manipulations were effective, allowing us to assess neural activity across two levels of sensitivity and two levels of decision criteria, in good shifters. The failure of some individuals to shift criteria despite explicit awareness is a known phenomenon (*36–42*), making poor shifters a useful control for analyses related to the decision criterion. Receiver operating characteristic (ROC) curves illustrate aggregated performance for good (Fig. 1G) and poor shifters (Fig. 1H), highlighting systematic differences between conditions, with points more spread out along both axes for good than for poor shifters. Reaction time plots reveal faster responses for high versus low confidence and correct versus incorrect trials across all test conditions (Fig. 1I), consistent with typical recognition memory performance (*34*). Additional behavioral (Fig. S1) and reaction time (Fig. S2) analyses are included in the Supplementary Materials.

### Neural recordings

We analyzed well-isolated single units with overall mean firing rates exceeding 0.3 Hz from 20 patients across 29 sessions (Supplementary Materials, table S1). Units meeting these criteria were recorded from the amygdala (*n* = 340), hippocampus (*n* = 355), dorsal anterior cingulate cortex (dACC; *n* = 195), and pre-supplementary motor area (pre-SMA; *n* = 310; Fig. 2A). We combined units from the amygdala and hippocampus (MTL; *n* = 695), and from the dACC and pre-SMA (MFC; *n* = 505). Assessments of neuron properties from this sample are included in the Supplementary Materials (Fig. S3).

**Fig. 2.**
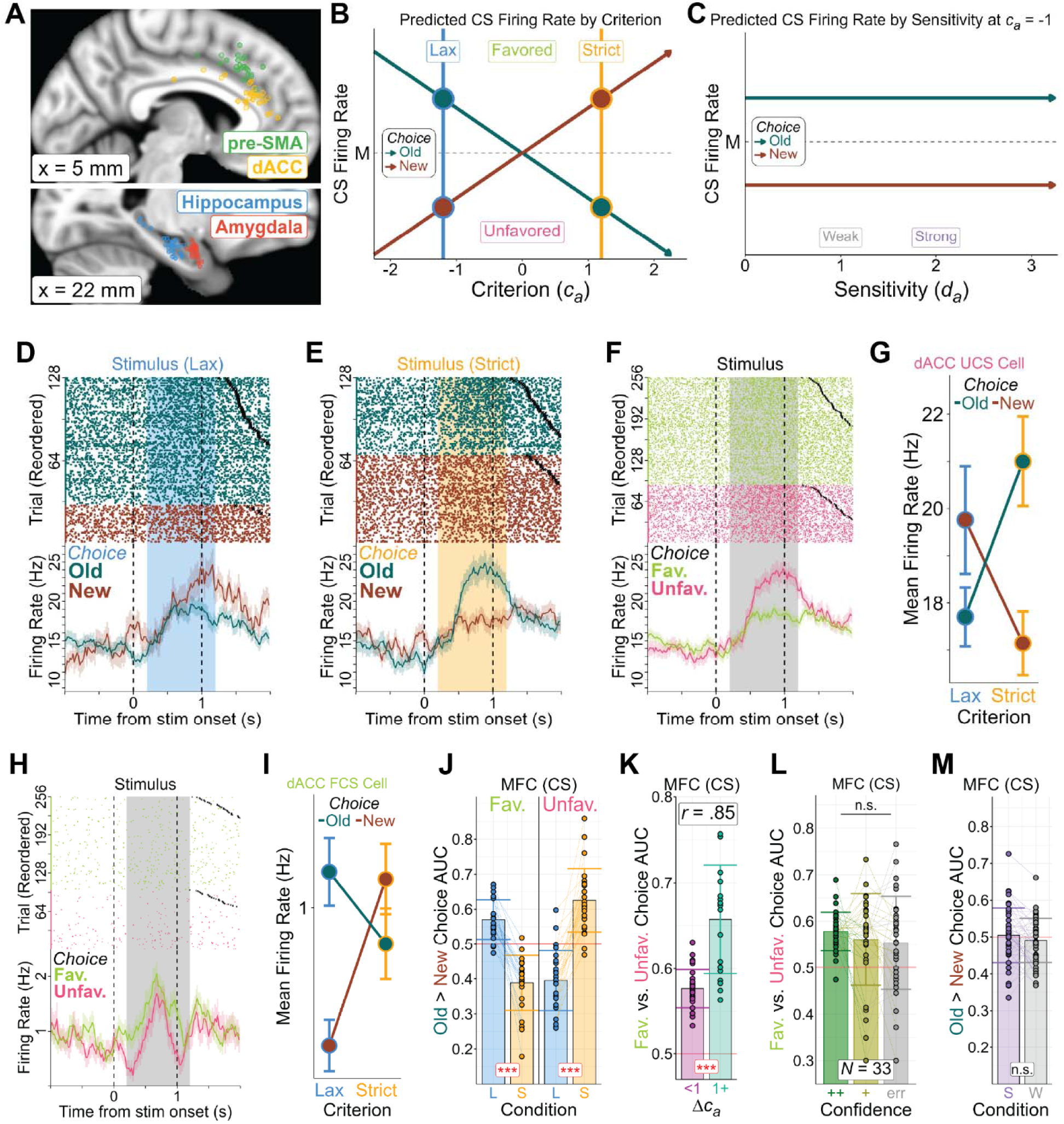
Electrode locations and criterion-selective (CS) neurons. (**A**) Electrode locations. Each dot marks the approximate location of a microwire bundle from which at least one neuron was recorded, merged onto a single MRI slice and plotted in Montreal Neurological Institute (MNI) 152 standard space. (**B**) Predicted CS neuron activity across decision criteria, indexed by *c_a_*. Hypothesized firing rate of a favored criterion-selective (FCS) neuron for “old” (green) and “new” (red) choices as the criterion shifts from lax (blue, negative *c_a_*) to strict (orange, positive *c_a_*). Larger criterion shifts are predicted to produce greater firing rate differences between favored and unfavored choices. Unfavored criterion-selective (UCS) neurons are predicted to show the opposite pattern (greater activity for unfavored than favored choices). (**C**) Predicted CS neuron activity across sensitivity, indexed by *d_a_*. Hypothesized firing rate of an FCS neuron for “old” and “new” choices at a fixed criterion (*c_a_* = −1), showing no change across sensitivity levels, reflecting independence from memory strength. (**D** to **G**) Example UCS dACC neuron. Raster plots (top) and peristimulus time histograms (PSTHs, bottom) show “old” and “new” choices under lax (**D**) and strict (**E**) criterion conditions. Trials are sorted by choice type and then by reaction time (fastest to slowest); black crosses (×) mark response time. The stimulus period used for analysis (200 to 1200 ms after stimulus onset) is highlighted in each raster and PSTH. (**F**) Trials regrouped into favored (lime; pooling lax old and strict new trials) and unfavored choices (pink; pooling lax new and strict old trials). (**G**) Mean ± SEM firing rates during the stimulus period show a significant choice-by-criterion interaction, with higher rates for “new” choices under a lax criterion and for “old” choices under a strict criterion. (**H** and **I**) Example FCS dACC neuron. Raster plot (top) and PSTH (bottom) show a preference for favored versus unfavored choices (**H**). Mean ± SEM firing rates likewise show a significant choice-by-criterion interaction (**I**). (**J**) Area under the curve (AUC) for “old” versus “new” choices in FCS (left) and UCS (right) MFC neurons under lax (blue, L) and strict (orange, S) criterion conditions (mean ± SD), showing significant differences across conditions. (**K**) AUC for favored versus unfavored choices from CS neurons in participants with smaller (Δ*c_a_* < 1; left) versus larger (Δ*c_a_* ≥ 1; right) criterion shifts (mean ± SD). AUC was strongly correlated with criterion shift magnitude [*r*(48) = .85], indicating that larger criterion shifts are associated with greater firing rate differences between favored and unfavored choices. (**L**) AUC for favored versus unfavored choices in CS MFC neurons, shown separately for high confidence correct (++, green), low confidence correct (+, yellow), and error (err, gray) trials (mean ± SD). The absence of significant differences is consistent with CS neuron firing being invariant to memory strength. (**M**) AUC for “old” versus “new” choices in CS MFC neurons under strong (purple, S) and weak (gray, W) sensitivity conditions (mean ± SD), showing no significant differences between conditions.

### Criterion-selective neurons in the MFC

We first tested whether neurons encode the current decision criterion, a population we refer to as “criterion-selective” (CS). We reasoned that CS neurons would show differential activity between the favored and unfavored choice during recognition tests, with the favored choice flipping between “old” and “new” decisions depending on whether the criterion is lax (favoring “old”) or strict (favoring “new”). The criterion would thus be reflected in the difference in firing rates between “old” and “new” choices, with larger differences associated with more extreme criteria and smaller differences with more neutral criteria. Furthermore, when comparing choices under a lax versus strict criterion, the difference in firing rate between “old” and “new” choices is expected to flip sign, reflecting a choice-by-criterion interaction (Fig. 2B). This prediction is motivated by prior fMRI findings showing that the old > new choice contrast in frontoparietal regions, including the MFC, is strongly modulated by criterion setting, suggesting that these regions flexibly encode the favored versus unfavored choice as a function of the adopted criterion (*27*, *32*). CS neurons were therefore identified as those exhibiting significantly different firing rates between favored choices (pooling lax old and strict new trials) and unfavored choices (pooling lax new and strict old trials) after stimulus onset, reflecting a specific interaction between the underlying criterion and choice type. We identified CS neurons in the MFC, but not the MTL (an example is shown in Fig. 2, D to G). In the MFC, the proportion of CS neurons was significantly greater in good shifters (11%, 45/400; *n* = 23 in dACC, *n* = 22 in pre-SMA) than in poor shifters (5%, 5/105; two-proportion z-test, *p* = .048). CS proportions in the MFC exceeded chance in good shifters (χ² test, *p* < .001) but not in poor shifters (*p* = .641), consistent with our prediction that CS neurons would be observed only in participants who successfully shift their criterion. In the MTL, CS neurons were not present in significant proportions (4%; 31/695; *p* = .767), regardless of whether individuals shifted their criteria or not.

We classified CS neurons into two subtypes: those firing more for favored choices (favored choice-selective, FCS) and those firing more for unfavored choices (unfavored choice-selective, UCS). This distinction was necessary because both subtypes were present in roughly equal proportions in the MFC (FCS: *n* = 24; UCS: *n* = 26), and collapsing across them would cancel out effects related to the criterion. Representative UCS and FCS neurons are shown in Fig. 2, D to I, with additional examples in the Supplementary Materials (Fig. S4).

For each CS neuron, we performed single-neuron ROC analyses within each criterion condition, computing the area under the curve (AUC) as the probability of correctly predicting “old” versus “new” choices from spike counts on individual test trials. AUC values above 0.5 indicate greater firing for “old” choices, values below 0.5 indicate greater firing for “new” choices, and 0.5 indicates chance. We adopted this convention for CS neurons specifically to illustrate how choice preferences change with the prevailing criterion. FCS neurons showed higher AUC in the lax (0.57 ± 0.06) than strict (0.39 ± 0.08) condition (*p* < .001), whereas UCS neurons showed the opposite pattern (lax, 0.40 ± 0.09; strict, 0.63 ± 0.09; *p* < .001; Fig. 2J). Thus, the activity of CS neurons differentiated between favored and unfavored choices regardless of whether the current choice was “old” or “new.”

We next assessed whether CS neuron firing tracks the magnitude of an individual’s criterion shift. For each CS neuron, we computed the AUC to assess the probability of correctly predicting favored versus unfavored choices based on single-trial spike counts (collapsed across criterion conditions), where 0.5 indicates no firing rate difference and larger values indicate greater differentiation. The AUC of CS neurons was strongly correlated with the magnitude of criterion shifting (Δ*c_a_*; *r*(48) = .85, *p* < .001), such that larger shifts were associated with greater differentiation between favored and unfavored choices. Accordingly, AUC was higher in CS neurons from participants with large criterion shifts (Δ*c_a_* ≥ 1) than small shifts (Δ*c_a_* < 1; *M* = 0.66 ± 0.06 versus 0.58 ± 0.02, *p* < .001; Fig. 2K), demonstrating that CS neuron activity tracks how extreme the criterion is.

If CS neurons encode the decision criterion as conceptualized in SDT, their activity should be insensitive to memory strength (see Fig. 2C). We tested this prediction by computing AUC values for favored versus unfavored choices across high confidence correct, low confidence correct, and incorrect trials, a method previously shown to index memory strength in MTL MS neurons through a monotonic decrease in AUC across these trial types (*34*). Among MFC CS neurons from participants who reliably used both confidence levels (*N* = 33), AUC did not differ significantly across these trial types [F(2,64) = 0.70, *p* = .498; Fig. 2L], indicating that CS neuron activity is invariant to memory strength, as SDT predicts. Similarly, AUC for “old” versus “new” choices, computed separately for the strong and weak memory strength conditions rather than for the criterion conditions, did not differ significantly between them (0.50 ± 0.07 versus 0.49 ± 0.06; *p* = .192), indicating that CS activity is also invariant to sensitivity (Fig. 2M). Together, these results indicate that, as predicted by SDT, CS neuron activity is insensitive to memory strength.

### Memory-selective neurons in the MFC and MTL

We next tested our second major prediction, that memory strength signals should be independent of the decision criterion (see Fig. 3A). To this end, we examined MS neuron activity as a function of the criterion condition. MS neurons comprised 16% of MFC neurons (80/505; *n* = 25 in dACC, *n* = 55 in pre-SMA) and 13% of MTL neurons (89/695; *n* = 37 in amygdala, *n* = 52 in hippocampus), both exceeding chance levels (χ² test of proportions, both *p* < .001). MS neuron proportions did not differ significantly between good and poor shifters in either the MFC (16% vs. 15%, *p* = .849) or MTL (13% vs. 12%, *p* = .634). Example MS neurons are shown in Fig. 3, C to H, and in the Supplementary Materials (Fig. S5).

**Fig. 3.**
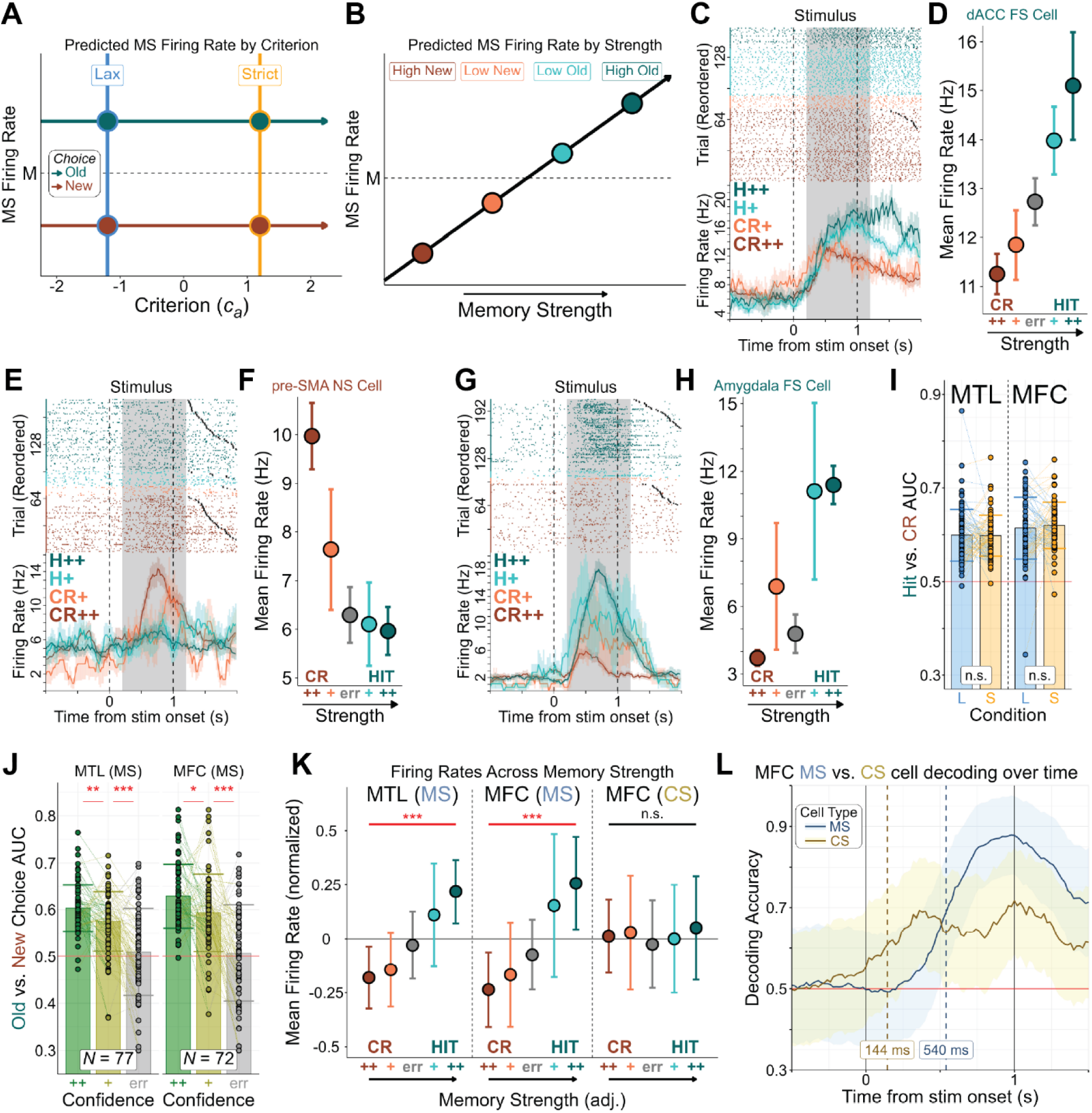
Memory-selective (MS) neurons. **(A)** Predicted MS neuron activity across decision criteria, indexed by *c_a_*. Hypothesized firing rate of a familiarity-selective (FS) neuron for “old” (green) and “new” (red) choices as the criterion shifts from lax (blue, negative *c_a_*) to strict (orange, positive *c_a_*). MS neurons are predicted to be unaffected by criterion shifts. (**B**) Predicted MS neuron activity across memory strength, indexed by confidence ratings. Hypothesized firing rate of an FS neuron as memory strength increases, with firing rate predicted to increase from high confidence “new” to high confidence “old” choices. (**C** and **D**) Example FS dACC neuron. (**C**) Raster plot (top) and peristimulus time histogram (PSTH, bottom) show high confidence hits (H++, green), low confidence hits (H+, cyan), low confidence correct rejections (CR+, coral), and high confidence correct rejections (CR++, red). Trials are sorted by choice type and then by reaction time (fastest to slowest); black crosses (×) mark response time. The stimulus period used for analysis (200 to 1200 ms after stimulus onset) is highlighted in each raster and PSTH. (**D**) Mean ± SEM firing rates during the stimulus period show a significant monotonic increase across memory strength, from CR++ to CR+ to errors (gray) to H+ to H++. (**E** and **F**) Example novelty-selective (NS) dACC neuron. Raster plot (top) and PSTH (bottom) show a preference for correct rejections over hits. (**F**) Mean ± SEM firing rates during the stimulus period show a significant monotonic decrease across memory strength. (**G** and **H**) Example FS amygdala neuron. (G) Raster plot (top) and PSTH (bottom) show a preference for hits over correct rejections. (**H**) Mean ± SEM firing rates during the stimulus period show a significant monotonic increase across memory strength. (**I**) Area under the curve (AUC) for hits versus correct rejections in MS MTL (left) and MFC (right) neurons under lax (blue, L) and strict (orange, S) criterion conditions (mean ± SD). No significant differences were observed across conditions, consistent with MS neuron firing being insensitive to the decision criterion. (**J**) AUC for hits versus correct rejections in MS MTL (left) and MFC (right) neurons, shown separately for high confidence correct (++, green), low confidence correct (+, yellow), and error (err, gray) trials (mean ± SD). AUC decreased significantly across trial types, indicating that MS neurons in both regions are sensitive to memory strength. (**K**) Normalized mean ± SD firing rates across choice types in MS MTL (left), MS MFC (middle), and CS MFC (right) neurons. MS neurons showed a monotonic increase in firing rate with memory strength, whereas CS neurons showed no such relationship. NS neurons showed a monotonic decrease in firing rate with memory strength; because values were calculated across NS and FS neurons together, NS values were inverted (adj.) to align with the direction of FS neurons for display. (**L**) Time-resolved single-trial population decoding accuracy for MFC CS neurons (favored versus unfavored choices) and MS neurons (hits versus correct rejections). CS neuron decoding reached half-maximum considerably earlier than MS neuron decoding (144 ms versus 540 ms after stimulus onset).

We classified MS neurons into two subtypes: those firing more on hit (H) trials (familiarity-selective, FS) and those firing more on correct rejection (CR) trials (novelty-selective, NS). For each MS neuron, we quantified memory strength encoding separately for each criterion condition using single-neuron ROC analysis, with AUC representing the probability of correctly discriminating H from CR trials based on single-trial spike counts (H > CR for FS; CR > H for NS). MS neurons in both regions showed comparable mean AUC under lax and strict conditions (MFC: 0.61 ± 0.07 vs. 0.62 ± 0.05, *p* = .522; MTL: 0.60 ± 0.06 vs. 0.60 ± 0.04, *p* = .896; Fig. 3I), consistent with MS neuron firing being insensitive to the decision criterion.

Among participants who reliably used both confidence levels, MTL MS neuron AUC (*N* = 77) decreased monotonically from correct high confidence (0.60 ± 0.05) to correct low confidence (0.58 ± 0.06; *p* = .002) to error trials (0.51 ± 0.09; *p* < .001; Fig. 3J, left), replicating previous findings (*34*). Extending these findings to MFC MS neurons (*N* = 72), AUC followed the same ordering: correct high confidence (0.63 ± 0.07) exceeded correct low confidence (0.59 ± 0.08; *p* = .012), which in turn exceeded error trials (0.51 ± 0.10; *p* < .001; Fig. 3J, right), demonstrating that memory strength modulates MS neuron activity in both regions. Accordingly, mean normalized firing rates increased monotonically with memory strength, as determined by Spearman correlations (ordered from high CR to low CR to errors to low H to high H, with NS neuron trial order reversed to maintain a consistent comparison direction), in both the MTL (ρ(76) = 0.69, 95% CI [0.63, 0.74], *p* < .001; Fig. 3K, left) and MFC (ρ(71) = 0.68, 95% CI [0.61, 0.74], *p* < .001; Fig. 3K, middle), confirming that MS neuron firing tracks memory strength across regions (compare with Fig. 3B). By contrast, MFC CS neurons showed no significant relationship (ρ(32) = 0.00, 95% CI [-0.18, 0.18], *p* = .973; Fig 3K, right), consistent with our prediction that these neurons are invariant to memory strength.

Finally, we compared the temporal dynamics of MFC CS and MS neuron populations using time-resolved single-trial population decoding. Decoding of favored versus unfavored choices in CS neurons reached half-maximum at 144 ms post-stimulus onset, considerably earlier than decoding of hits versus correct rejections in MS neurons (540 ms; Fig. 3L), demonstrating that CS neurons signal the decision criterion well before mnemonic information becomes available. Further discussion of the potential temporal relationship between CS and MS neurons appears in the Supplementary Materials (Fig. S4A).

### Visually-selective neurons in the MFC and MTL

Visually-selective neurons contribute to declarative memory by representing specific image categories rather than memories for individual items (*16*, *34*), and should therefore be unaffected by criterion shifts (see Fig. 4A) or changes in memory strength (see Fig. 4B). Since our task included four image categories evenly distributed across conditions and old versus new image types, we examined how VS neuron activity was modulated by criterion shifts and memory strength.

**Fig. 4.**
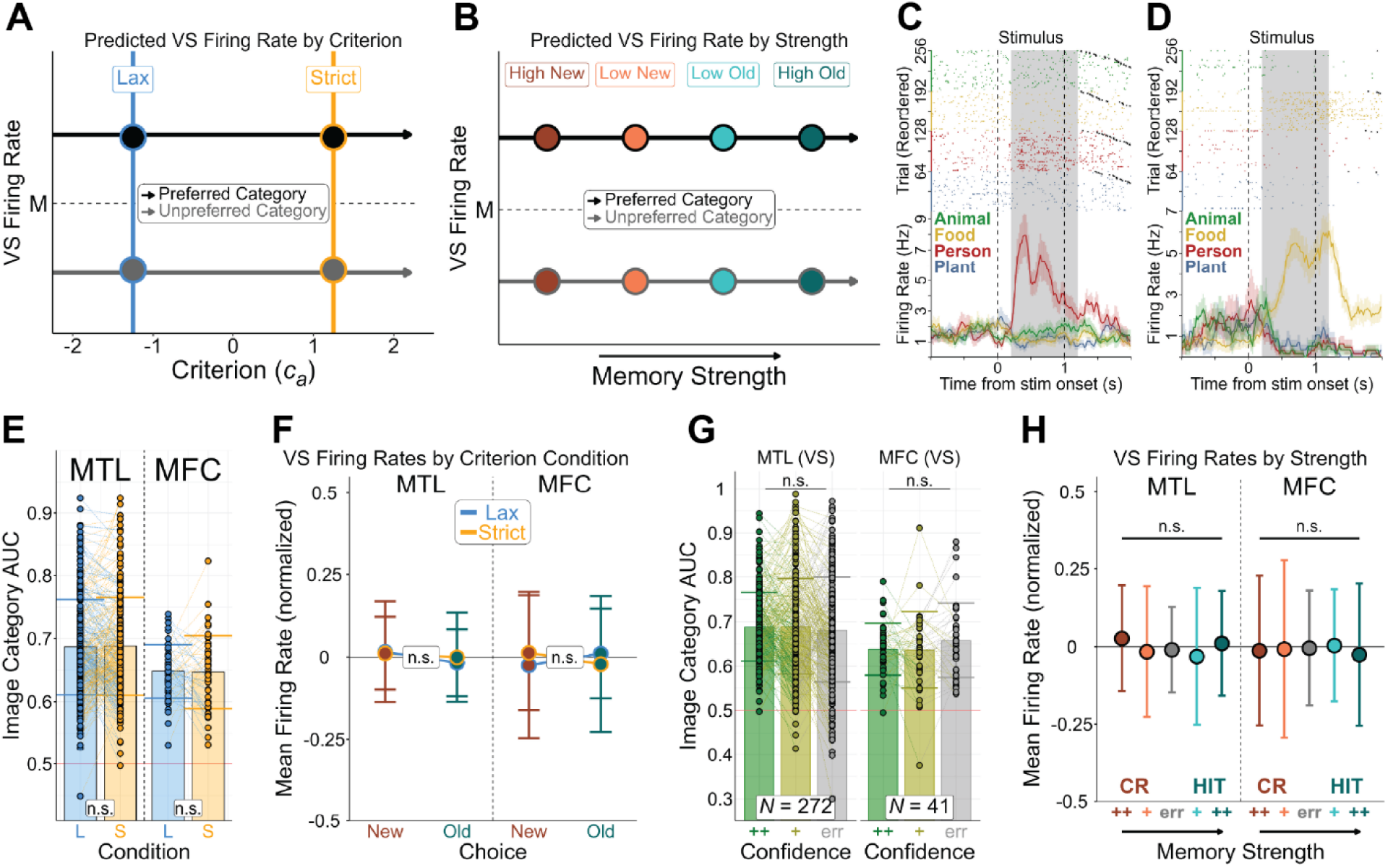
Visually-selective (VS) neurons. (**A**) Predicted VS neuron activity across decision criteria, indexed by *c_a_*. Hypothesized firing rate for the preferred image category (black) and unpreferred image category (gray) as the criterion shifts from lax (blue, negative *c_a_*) to strict (orange, positive *c_a_*). VS neurons are predicted to be unaffected by criterion shifts. (**B**) Predicted VS neuron activity across memory strength, indexed by confidence ratings. VS neurons are predicted to be unaffected by memory strength. (**C**) Example VS hippocampus neuron showing a preference for images of people. Raster plot (top) and peristimulus time histogram (PSTH, bottom) show trials in which the image category was of an animal (green), food (yellow), a person (red), or a plant (blue). Trials are sorted by image category and then by reaction time (fastest to slowest); black crosses (×) mark response time. The stimulus period used for analysis (200 to 1200 ms after stimulus onset) is highlighted in each raster and PSTH. (**D**) Example VS dACC neuron showing a preference for images of food. (**E**) Area under the curve (AUC) for the preferred versus unpreferred image category in VS MTL (left) and MFC (right) neurons under lax (blue, L) and strict (orange, S) criterion conditions (mean ± SD). No significant differences were observed across conditions, consistent with VS neurons being insensitive to the decision criterion. (**F**) Mean normalized firing rates in VS MTL (left) and MFC (right) neurons show no significant differences across choice types and criterion conditions. (**G**) AUC for preferred versus unpreferred image category in VS MTL (left) and MFC (right) neurons, shown separately for high confidence correct (++, green), low confidence correct (+, yellow), and error (err, gray) trials (mean ± SD). The absence of significant differences is consistent with VS neuron firing being invariant to memory strength. (**H**) Normalized mean ± SD firing rates across choice types in VS MTL (left) and VS MFC (right) neurons. VS neurons show no significant modulation across memory strength.

VS neurons comprised 10% of MFC neurons (50/505; *n* = 24 in dACC, *n* = 26 in pre-SMA) and 44% of MTL neurons (308/695; *n* = 162 in amygdala, *n* = 146 in hippocampus), with both proportions exceeding chance (χ² test of proportions, both *p* < .001). VS neuron proportions did not differ significantly between good and poor shifters in either the MFC (11% vs. 8%, *p* = .379) or MTL (45% vs. 44%, *p* = .776). Representative examples of VS neurons are shown in Fig. 4, C and D, and in the Supplementary Materials (Fig. S6).

For each VS neuron, we conducted single-neuron ROC analyses across criterion conditions, calculating the AUC as the probability of predicting the preferred stimulus category (highest firing rate) versus the least preferred category (lowest firing rate) from spike counts on individual trials. VS neurons in both regions showed comparable AUC under lax and strict criterion conditions (MTL: 0.69 ± 0.08 versus 0.69 ± 0.08, *p* = .721; MFC: 0.65 ± 0.04 versus. 0.65 ± 0.06, *p* = .902; Fig. 4E), and normalized firing rates did not differ between favored and unfavored choices in either region (MTL: *M* = 0.01, 95% CI [−0.00, 0.03], *p* = .181; MFC: *M* = 0.03, 95% CI [−0.02, 0.08], *p* = .169; Fig. 4F). This result is consistent with our prediction that VS neuron firing is insensitive to the decision criterion (compare with Fig. 4A).

Among participants who reliably used both confidence levels, mean AUC did not significantly differ across correct high confidence, correct low confidence, and error trials in either the MTL (*N* = 272; F(2,542) = 0.62, *p* = .540; Fig. 4H) or MFC (*N* = 41; F(2,80) = 0.99, *p* = .375; Fig. 4G), and normalized firing rates showed no relationship with memory strength in either region (MTL: ρ(264) = −0.05, 95% CI [−0.11, 0.01], *p* = .127; MFC: ρ(40) = 0.00, 95% CI [−0.16, 0.16], *p* = .975; Fig. 4H). Together, these results indicate that VS neurons encode stimulus category independently of memory strength (compare with Fig. 4B).

### Baseline encoding of the criterion condition in the MFC

In addition to characterizing neurons based on firing rate changes after stimulus onset, we examined whether the criterion condition was reflected in baseline firing rates before test image onset. Since conditions were blocked and participants were informed of the associated error penalties at the start of each block, they could maintain awareness of the criterion condition throughout the block. Based on prior single-unit findings implicating the MFC in task context maintenance (*16*, *18*, *43*), we predicted that criterion condition encoding during the baseline would be preferentially represented in the MFC. In the MFC, 13% of neurons (67/505; *n* = 23 in dACC, *n* = 44 in pre-SMA; χ² test of proportions, *p* < .001) exhibited significant baseline firing rate differences between lax and strict criterion conditions, compared with only 8% of MTL neurons (58/695; *n* = 30 in amygdala, *n* = 28 in hippocampus; *p* < .001). The MFC proportion significantly exceeded that of the MTL (*p* = .006). The proportion of MFC neurons encoding criterion condition was significant in both good (12%; 48/400; *p* < .001) and poor (18%; 19/105; *p* < .001) shifters, suggesting that a failure to shift criteria does not reflect an inability to represent and remember the current criterion condition. Example baseline criterion condition neurons from a good shifter (Fig. 5, A to C) and a poor shifter (Fig. 5, D to F) are shown.

**Fig. 5.**
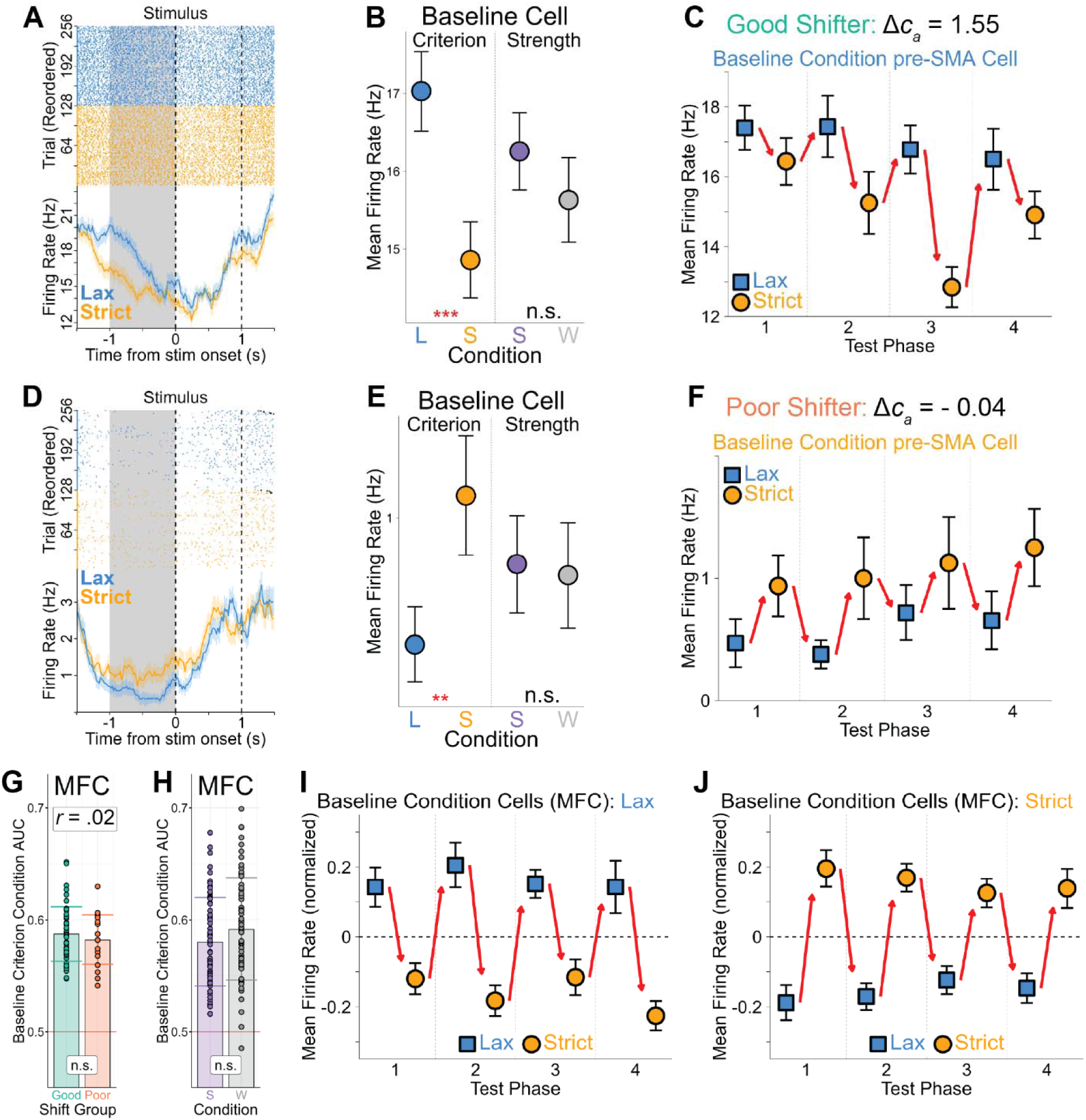
Baseline criterion condition neurons do not predict criterion placement. (**A** to **C**) Example baseline criterion condition pre-SMA neuron from a good shifter, favoring the lax over the strict criterion condition. (**A**) Raster plot (top) and peristimulus time histogram (PSTH, bottom) show trials in the lax (blue) and strict (orange) criterion conditions. Trials are sorted by criterion condition and then by reaction time (fastest to slowest); black crosses (×) mark response time. The baseline period used for analysis (−1000 to 0 ms before stimulus onset) is highlighted in each raster and PSTH. (**B**) Mean ± SEM firing rates during the baseline period in the lax (blue, L) and strict (orange, S) criterion conditions and in the strong (purple, S) and weak (gray, W) memory strength conditions, showing a significant difference between criterion conditions but not between memory strength conditions. (**C**) Mean baseline firing rates across criterion conditions, separately for each test phase, showing greater firing in the lax condition across all four test phases. (**D** to **F**) Example baseline criterion condition pre-SMA neuron from a poor shifter, favoring the strict over the lax criterion condition. (**D**) Raster plot (top) and PSTH (bottom) show trials in each criterion condition. (**E**) Mean ± SEM firing rates during the baseline period show a significant difference between criterion conditions but not between memory strength conditions. (**F**) Mean baseline firing rates across criterion conditions are consistently higher in the strict than the lax condition across test phases. (**G**) Baseline criterion condition AUC did not differ significantly between the good (cyan) and poor (coral) shift groups, consistent with these neurons not being diagnostic of criterion placement. (**H**) Baseline criterion condition AUC computed separately for the strong (purple, S) and weak (gray, W) memory strength conditions showed no significant difference, consistent with these neurons being specifically attuned to the criterion condition rather than the memory strength condition. (**I** and **J**) Mean normalized firing rates across all MFC baseline condition neurons preferring the lax (**I**) and strict (**J**) criterion conditions, demonstrating sustained preferential firing for the preferred criterion condition across the entire testing session.

For each MFC neuron encoding the criterion condition at baseline, we performed single-neuron ROC analysis, computing AUC as the probability of correctly predicting the current criterion condition from spike counts on individual trials. AUC exceeded chance in both good (0.59 ± 0.02) and poor (0.58 ± 0.02) shifters (*p* < .001 for both), with no significant difference between groups (*p* = 0.389; Fig. 5G). Unlike CS neurons, which are strongly associated with criterion shifting, these baseline criterion condition neurons showed no correlation with the extent of individual criterion shifting (Δ*c_a_*; *r*(65) = .02, *p* = .867), suggesting that criterion condition encoding at baseline is unrelated to criterion shifting behavior. We also computed AUC separately for each memory strength condition and found no significant difference between strong and weak conditions (0.58 ± 0.04 versus 0.59 ± 0.05, *p* = .199; Fig 5H), consistent with these neurons encoding the criterion condition independently of memory strength. Normalized firing rates across all MFC baseline condition neurons, averaged within each of the four test phases, are shown separately for neurons preferring the lax (Fig. 5I) and strict (Fig. 5J) criterion conditions, demonstrating that this population maintains preferential firing for its preferred criterion condition throughout the testing session. Together, these results indicate that a subpopulation of MFC neurons encodes the criterion condition during the baseline period, independently of whether participants shift their criterion. This suggests that these neurons represent the context of the current criterion rather than diagnostic information about its precise placement.

### Decision-making model

We implemented a computational model to examine how CS neurons influence and improve memory-based decisions. We extended the “balance of evidence” race model previously used to explain how MTL MS neurons encode accuracy and confidence during recognition judgments (*34*). In this “memory strength” model, the firing rate difference between a pair of NS and FS neurons is integrated over time (Fig. 6A). At trial end, an “old” choice is made if the accumulated evidence (EV) is positive, and a “new” choice is made if EV is negative.

**Fig. 6.**
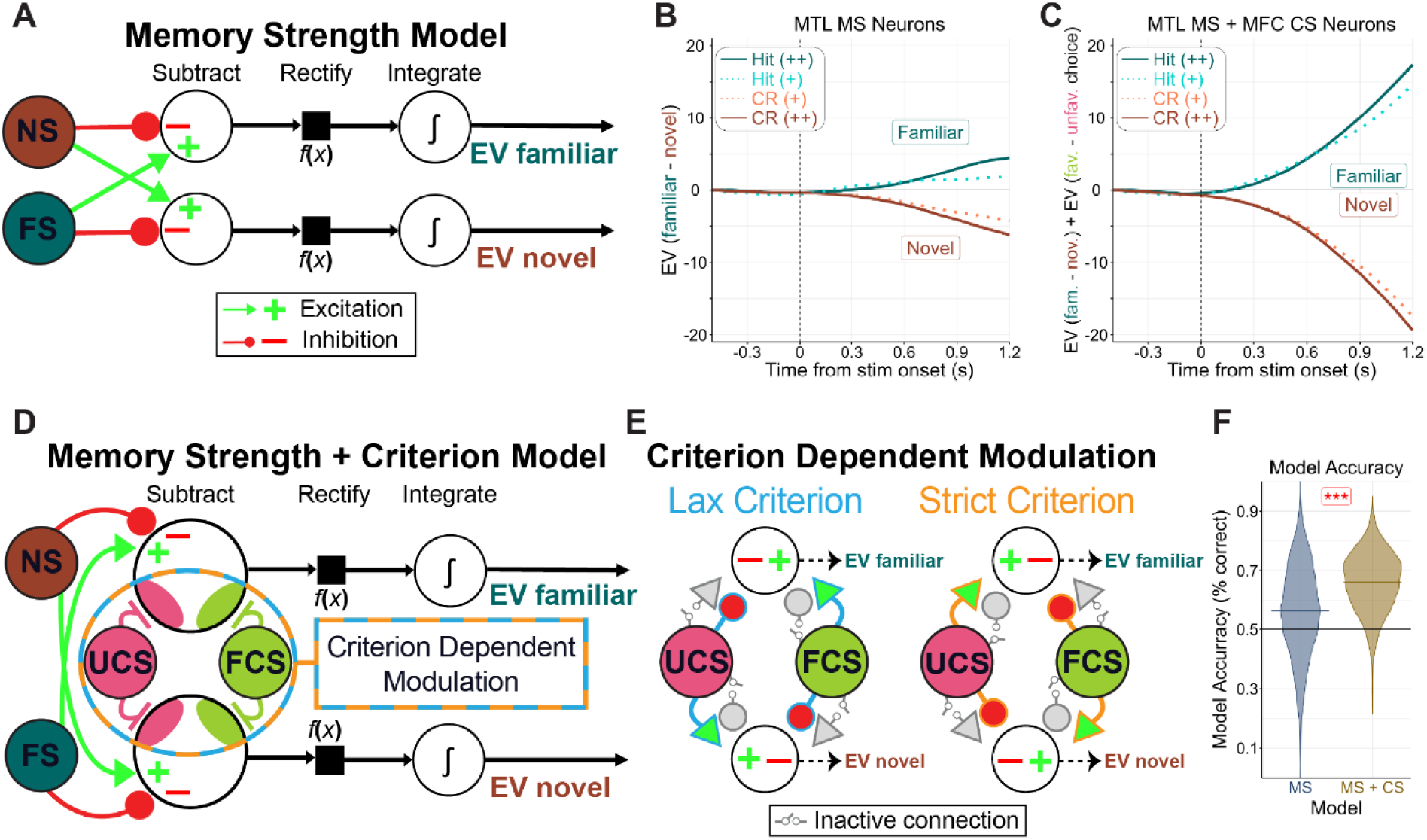
Decision-making models. (**A**) Circuit diagram of the memory strength model, which accumulates the firing rate difference between a pair of NS and FS neurons over time. Green arrows and “+” signs indicate excitation, whereas red circles and “−” signs indicate inhibition. (**B**) Model output for all MTL NS-FS pairs (*n* = 720) for high (++, solid lines) and low (+, dashed lines) confidence hits (dark green) and correct rejections (CRs; dark red), showing greater separation for high than low confidence choices. (**C**) Model output for all MTL NS-FS and MFC UCS-FCS quartets (*n* = 449,280) for high and low confidence hits and CRs, combined across criterion conditions, showing faster evidence accumulation for hits and correct rejections than the memory strength model alone (**D**) Circuit diagram of the memory strength plus criterion model, which integrates evidence from paired NS-FS and UCS-FCS neurons jointly to model the decision criterion along with memory strength. (**E**) Circuit diagram of UCS-FCS neurons, shown separately for the lax (blue) and strict (orange) criterion conditions. Depending on the criterion, one set of connections is inactive (gray) while the other independently modulates EV over time alongside the NS-FS pairs. (**F**) Model accuracy at trial end for the memory strength model alone (blue, left) and the memory strength plus criterion model (yellow, right); accuracy was significantly higher when CS neurons were incorporated into the model.

We extended this model to incorporate CS neurons by adding pairs of UCS and FCS neurons as additional inputs, with the evidence integration summing the pooled input from both MS and CS neurons (memory strength + criterion model; Fig. 6D). Unlike MS neurons, which maintain their connectivity regardless of the criterion, the connectivity formed by the UCS and FCS neurons and the summation unit changes as a function of the criterion setting. Changing the criterion, in this model, is therefore implemented by changing connectivity (Fig. 6E). In effect, switching between the two criterion levels introduces a decision bias, whereby EV accumulates faster for familiarity under a lax criterion but faster for novelty under a strict criterion.

We next assessed how well the original and extended models performed when fed the recorded data as an input. We first validated the memory strength model, before incorporating CS neurons, across all 720 MTL NS-FS pairs from participants with sufficient trials of each confidence rating. At trial end (1200 ms after stimulus onset), the model successfully distinguished hits from correct rejections on individual trials (*p* < .001), and accuracy was significantly higher for high than low confidence judgments (58.5% ± 15.0% vs. 54.1% ± 16.3%; *p* < .001 ; Fig. 6, B and F).

We then incorporated CS neurons by pairing each MTL NS-FS pair with each MFC UCS-FCS pair, yielding 449,280 quartets. The combined model substantially improved accuracy (66% ± 10%; Fig. 6, C and F) relative to the memory strength model alone (56% ± 16%; *p* < .001). Together, these results indicate that recognition memory judgments predicted by the model improve when they rely on the joint activity of MS and CS neurons, with MS neurons tracking memory strength and CS neurons encoding the decision criterion.

## Discussion

We dissociated neural activity underlying the two major hypothesized components of memory-based decision making: the decision criterion and memory strength. Our central finding is the identification of neurons in the MFC, but not the MTL, that track whether an individual has adopted a lax or strict decision criterion. These “criterion-selective” (CS) neurons were present only in individuals who demonstrably shifted their decision criterion behaviorally. The magnitude of behavioral criterion shifts scaled with the degree of differential firing in CS neurons, suggesting that CS neuron activity in the MFC is behaviorally relevant for criterion shifting. As predicted by SDT, CS neuron activity was insensitive to memory strength, indicating that these neurons track the decision criterion independently of the underlying memory signal. CS neurons also signaled the favored versus unfavored choice well before MS neurons signaled memory strength, suggesting that CS neurons influence memory-based decisions earlier than MS neurons. MS neurons, on the other hand, showed the opposite selectivity profile: their activity scaled with confidence and accuracy, indexing memory strength, but was unaffected by the decision criterion. Together, these results reveal two functionally distinct neural populations in the MFC that selectively and independently encode the decision criterion and memory strength during recognition memory.

Our approach of identifying neural populations that independently represent the decision criterion from the underlying signal complements prior macaque research that has identified populations that integrate these processes into a single decision variable (*7–12*). Since the decision criterion and signal processing are computationally independent (*6*), characterizing the neural mechanisms of each on its own is a necessary step toward understanding how these signals are ultimately integrated to produce a decision. To model how these independent neural populations jointly predict decisions, we built on a prior model that predicted behavioral responses from the neural readout of competing novelty-selective and familiarity-selective MTL neuron pairs (*34*). The original model lacked a mechanism for the decision criterion, so we extended it by incorporating MFC CS neurons, operationalized as the readout of competing unfavored choice-selective and favored choice-selective neuron pairs. Incorporating CS neurons substantially improved prediction accuracy, demonstrating that integrating activity from both CS and MS neurons predicts participants’ memory-based decisions better than MS activity alone.

Beyond characterizing CS and MS neurons, we also examined “visually-selective” (VS) neurons previously identified in the MFC (*16*) and MTL (*16*, *34*), and found that VS neuron activity is independent of both the decision criterion and memory strength in both regions. Our results reveal three functionally dissociable neural populations across the MFC and MTL: CS neurons that selectively encode the decision criterion (MFC only), MS neurons that selectively encode memory strength, and VS neurons that encode stimulus category independently of both, each serving a distinct computational role in the formation of recognition judgments.

We also identified neurons that encoded the criterion condition during baseline periods. Baseline criterion condition representations were stronger in the MFC than the MTL, suggesting that the MFC plays a more prominent role in maintaining the context of the criterion condition during memory tests, consistent with prior reports implicating the MFC in task context representations (*16*, *18*, *43*). Similar differences in baseline neural activity between conditions designed to induce criterion shifts have been reported in macaques (*9–12*) and mice (*44*, *45*). However, the presence of a baseline signal does not by itself establish that it represents the decision criterion. In our data, this baseline signal was present even in individuals who failed to shift their criterion, and showed no relationship with the magnitude of that shift. This indicates that these neurons track context rather than the criterion per se, and that criterion inflexibility reflects an unwillingness to shift rather than a failure to represent and remember the current criterion condition (*36–42*). Differentiating signals that track decisional context from those that track the criterion itself is therefore essential to understanding how decision biases inform judgments.

Our findings help resolve a longstanding debate over whether and how the decision criterion is distinctly represented from memory strength in the brain (*14*, *20–33*). Although the MFC is well known to represent memory-based decisions (*20*), with an identified cellular correlate (*16*), a cellular correlate of the decision criterion itself had not been identified. Previous fMRI work demonstrated that the MFC represents the decision criterion (*27*, *32*), but its limited spatial resolution left it unclear whether and how the criterion and memory strength were represented distinctly, motivating the current study. Using single-neuron recordings, we identified two functionally distinct MFC populations, CS neurons and MS neurons, whose activity scaled with the extremity of the decision criterion and the strength of memory, respectively. Inconsistent fMRI findings may therefore reflect an inability to resolve these intermingled populations, compounded by cross-study variability in criterion placement and memory strength. Tasks that elicit extreme criteria and weak memories may preferentially engage CS over MS neurons, favoring a criterion account, whereas tasks inducing strong memories and more neutral criteria may do the opposite, favoring a memory strength account. In reality, both populations coexist in the MFC and jointly contribute to the recognition judgment, consistent with the predictions of SDT. More broadly, these findings show that the decision criterion and the evidence it acts on can be represented by distinct neural populations, and that understanding how each process is represented independently of the other is a necessary step toward understanding individual behavior and how such signals are ultimately integrated to form a decision.

## Materials and methods

### Subjects

Epilepsy patients undergoing seizure monitoring (36 sessions from 24 subjects) participated in the experiment from their hospital beds using a laptop while Behnke-Fried depth electrodes recorded single neurons. Seven sessions from four patients were excluded (six due to chance performance in both the weak and strong memory conditions, and one due to the absence of usable units in the MFC or MTL) leaving a final sample of 29 sessions from 20 subjects for analysis. Eighteen patients (22 sessions) were recorded at Cedars-Sinai Medical Center, and six (14 sessions) were recorded at Toronto Western Hospital. All protocols were approved by the institutional review boards of Cedars-Sinai Medical Center, the California Institute of Technology, and Toronto Western Hospital.

### Stimuli

Stimuli were drawn from the LaMem dataset (*46*) across four categories (animals, food, people, and plants) and center-cropped to 400×400 pixels. For each image, a single memorability score was computed by averaging its scores across all test, training, and validation sets. We selected images with mean memorability scores between 0.60 and 0.92 and distributed them evenly across categories to control overall memorability. The task comprised three versions to accommodate multiple testing sessions, each containing 256 unique stimuli evenly distributed across the four categories.

### Task

The recognition memory task used a fully crossed 2×2 factorial design that manipulated memory strength (strong versus weak) and the decision criterion (lax versus strict; Fig. 1B). During each of the four study phases, participants viewed 32 unique images, shown either once (weak) or twice (strong), yielding 48 presentations (Fig. 1C). Images were presented for 1000 ms, preceded by a jittered fixation crosshair (800–1200 ms), with yes/no attention check questions appearing after 8 randomly selected images asking participants to identify whether the preceding image belonged to a specific category (e.g., food or a person). At test, participants earned 4 points for a correct choice, lost 8 points for critical errors, and were not penalized for noncritical errors. Critical errors alternated pseudorandomly between incorrect “old” (strict) and incorrect “new” (lax) choices, and participants were always informed of the point structure. Following each study phase, participants completed four test blocks corresponding to the four test condition combinations (lax strong, lax weak, strict strong, strict weak). Each test block consisted of 16 trials (8 old, 8 new) presented in a randomized order (Fig. 1D). Strong memory strength blocks included old images presented twice during study, whereas weak blocks included old images presented once. Trials began with a jittered crosshair (1000–1500 ms), followed by a 1000 ms test image presentation, and concluded with a response screen where participants simultaneously indicated “old” or “new” and rated confidence as “high” or “low.” Stimuli consisted of four visual categories (animals, food, people, and plants), evenly distributed across conditions and item types. After each test block, subjects received feedback displaying their total points earned so far and the percentage of the maximum possible score for that block (out of 64). The experiment included 16 test blocks (4 per condition combination), totaling 256 test trials. Prior to the task, participants read instructions and completed practice trials to ensure comprehension of the task and point system, so that any failure to shift criteria could not be attributed to a misunderstanding of task conditions.

### Behavioral Analysis

Recognition memory performance was assessed using ROC analyses implemented in the ROC Toolbox (*47*), which models memory strength as a latent decision variable. “New” items followed a standard normal distribution, while “old” items followed a normal distribution with distinct mean (*μ*) and standard deviation (*σ*). Three thresholds (*t₁*, *t₂*, *t₃*) defined the categorical boundaries among the four response types within each condition (old and new with high or low confidence). Distributional parameters were held constant across criterion conditions, and threshold parameters were held constant across difficulty conditions, yielding 10 free parameters (2 *μ*, 2 *σ*, 6 *t*). Discriminability (*d_a_*) was computed from these parameters as follows:

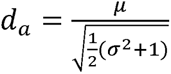

Criterion placement (*c_a_*) was computed by:

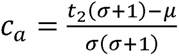

Criterion shifting (Δ*c_a_*) was computed by:

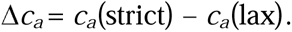

Differences in these parameters between conditions were assessed using two-tailed *t*-tests across participants, with significance determined by *p* < .05. Aggregated ROC curves (Fig. 1, G and H) were computed by summing all response categories across participants, then performing ROC analyses on the group sum.

Reaction time was defined as the interval from stimulus onset to the participant’s response. Responses could only be made after the test image disappeared, 1000 ms after stimulus onset. Pearson correlations were used to assess relationships between criterion shifting measures (Δ*c_a_*) and neural AUC measures, whereas Spearman correlations were used to assess the rank order of firing rates across trial types hypothesized to track memory strength according to SDT (high new < low new < errors < low old < high old) (*35*). Correlations were considered significant at *p* < .05.

### Electrophysiology

Bilateral recordings from the amygdala, hippocampus, dACC, and pre-SMA were obtained using Behnke–Fried hybrid depth electrodes with eight microwires. Each microwire recorded extracellular signals filtered from 0.1 to 9,000 Hz at a sampling rate of 32 kHz (ATLAS system, Neuralynx Inc.). One microwire per depth electrode served as a local reference, and electrode locations were verified using post-operative MRI and CT scans.

### Spike detection and sorting

Raw signals were bandpass filtered between 300 and 3,000 Hz using a zero-phase lag filter. Spikes were detected and sorted with the semi-automated OSort algorithm (*48*), and only well-isolated units verified through manual inspection were included in subsequent analyses. Peristimulus time histogram (PSTH) plots depicted below each raster plot were computed using a 200-ms sliding window with a step size of 20 ms.

### Selection of units

Selective neurons were identified using a bootstrap ANOVA with 5,000 iterations (*p* < .05) applied to firing rates measured 200 to 1200 ms after test stimulus onset, or 1000 to 0 ms before stimulus onset for neurons tracking the criterion condition during baseline. CS neurons were defined by significantly different firing rates between favored (lax old, strict new) and unfavored (lax new, strict old) choices, with those firing more strongly for favored choices classified as FCS and the remainder as UCS. MS neurons were defined by significant firing rate differences between hit and correct rejection trials, with neurons exhibiting higher firing rates for hits classified as FS and the remainder as NS. VS neurons were defined by significant firing rate differences across the four image categories, with selectivity assigned to the category eliciting the highest firing rate. Neurons encoding the criterion condition during baseline were defined by a significant firing rate difference between lax and strict conditions, using firing rates measured 1000 to 0 ms before test stimulus onset.

### Single-neuron ROC analysis

ROC curves were constructed from spike counts in the 200 to 1200 ms window after test stimulus onset for CS, MS, and VS neurons, and from spike counts in the −1000 to 0 ms window before test stimulus onset for neurons encoding the criterion condition during baseline. Detection thresholds spanned the full range of unique spike counts observed for each neuron, over which the AUC was computed by integrating the ROC curve. AUC was computed separately for each criterion (lax versus strict) condition. To maintain consistent comparisons, the first group in each contrast included only within-condition trials, while the second group was pooled across conditions (e.g., lax hits vs. all correct rejections), unless otherwise specified. Each comparison required a minimum of three trials per response type within each condition.

For CS neurons, ROCs contrasted old versus new choices to assess how choice type preference varied across criterion conditions. Since CS neurons reverse their choice type preference across criterion conditions, both old and new choices were analyzed within-condition to preserve this effect (e.g., strict old choices vs. strict new choices). For MS neurons, ROCs contrasted hits versus correct rejections to assess each neuron’s preferred response type, with correct rejection trials pooled across conditions for FS neurons and hit trials pooled across conditions for NS neurons. For VS neurons, ROCs compared trials from the image category eliciting the highest firing rate against those from the category eliciting the lowest, with the latter pooled across conditions. For confidence analyses, AUC was computed separately for correct high confidence, correct low confidence, and error trials, with the first group separated by confidence level and the second group pooling trials across both confidence levels (e.g. high confidence hits versus all correct rejections). For neurons encoding the criterion condition during baseline, AUC was computed between lax and strict conditions, either together or separately for the strong and weak memory strength conditions. Statistical comparisons between AUC values were performed using two-tailed *t*-tests or ANOVAs, with *p* < .05 as the significance threshold.

### Population decoding

MFC CS and MS neurons were each pooled separately across sessions to form two distinct pseudo-populations for time-resolved single-trial decoding. The CS pseudo-population was assessed for its ability to discriminate favored from unfavored choices, while the MS pseudo-population was assessed for its ability to discriminate hits from correct rejections. Firing rates were computed using a 400 ms sliding window advanced in 20 ms steps over a −500 to 1500 ms peri-stimulus window and z-score normalized. A linear support vector machine (SVM) with 10-fold cross-validation was used to estimate decoding performance. To balance trial counts across conditions, 10 trials were randomly sampled from each condition per iteration, and this procedure was repeated for 10,000 iterations; means and standard deviations at each time point are reported across iterations. Decoding latency was defined as the time at which decoding accuracy reached half its maximum value above chance (0.5), estimated via linear interpolation between consecutive time points.

### Decision Making Model

The decision-making model extends a previously described “balance of evidence” race model that predicted recognition accuracy and confidence from the competing activity of MTL NS and FS neurons (*34*). We applied this model (the memory strength model; Fig. 6A) to our dataset by inputting spiking activity 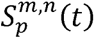 from each MTL NS (*i*) and FS (*j*) neuron pair for each correct trial (*k*) and integrating the difference in activity over time 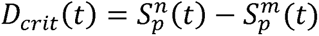. Spikes were binned into 250 ms windows with 100 ms steps, spanning from -500 ms to 1,200 ms relative to test stimulus onset, and firing rates were z-score normalized to each neuron’s mean firing rate across the entire session. Evidence for familiarity *EV*_fam_(*t*) and novelty *EV*_nov_(*t*) accumulated over time as follows:

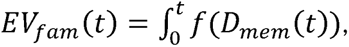

and

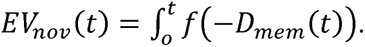

An “old” decision was made if the balance of evidence Δ*E*_mem_(*t*) = *EV*_fam_(*t*) – *EV*_nov_(*t*) was positive at 1,200 ms after stimulus onset, and a “new” decision if negative. The model was run on all possible MTL NS-FS pairs (*n* = 720) with at least 3 correct trials in each of the four response types (old and new, each at high or low confidence).

We extended this framework to incorporate the decision criterion (the memory strength + criterion model; Fig. 6, D and E) by additionally inputting spiking activity 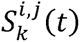 from each MFC UCS (*m*) and FCS (*n*) neuron pair for each trial (*p*) and integrating the difference in activity over time 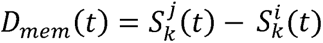. As with MS neurons, spikes were binned into 250 ms windows with 100 ms steps spanning -500 ms to 1,200 ms relative to stimulus onset, with firing rates z-score normalized to each neuron’s mean across the session. Evidence for favored *EV*_fav_(*t*) and unfavored *EV*_unfav_(*t*) choices accumulated over time as follows:

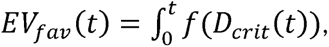

and

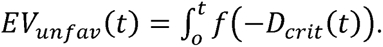

We paired all MTL NS-FS pairs with all MFC UCS-FCS pairs, yielding 449,280 quartets. For each quartet, the combined “memory strength and criterion” balance of evidence was computed as Δ*E_mem+crit_*(*t*) = *EV*_fam_(*t*) – *EV*_nov_(*t*) + *EV*_fav_(*t*) – *EV*_unfav_(*t*), jointly capturing memory strength, via MTL MS neurons, and the decision criterion, via MFC CS neurons. To maintain consistency in the direction of the criterion across neurons, trials were compared separately for the lax and strict criterion conditions, such that *EV*_fav_(*t*) represented old choices in the lax condition and new choices in the strict condition, with the reverse holding for *EV*_unfav_(*t*).

## ACKNOWLEDGMENTS

We thank the patients and their families for their generous participation in this research. We further thank the staff of the Epilepsy Monitoring Unit and the Biomedical Imaging Research Institute at Cedars-Sinai Medical Center, as well as the staff at Toronto Western Hospital, for patient care and research support. We also thank members of the Rutishauser and Valiente labs for their assistance with data collection, analyses, and feedback on this work. We thank B. Layher for assistance with data analysis and figure preparation. **Funding:** This work was supported by the NIH BRAIN Initiative through the NIH Office of the Director (U01NS117839 to U.R.). **Author contributions:** E.L., M.B.M., and U.R. designed the study. E.L. and I.S. performed the experiments. E.L. analyzed the data. C.M.R., J.M.C., and L.M.B. provided patient care and facilitated experiments. T.A.V. and A.N.M. performed surgery and supervised clinical work. E.L., M.B.M., and U.R. wrote the paper with input from all authors. **Competing interests:** The authors declare no competing interests. **Data and materials availability:** Data and code to reproduce the main results will be made publicly available on OSF upon publication.

