## Supplemental Materials for "Criterion-selective neurons in the human medial frontal cortex track decision thresholds during memory-based decision making"

**Materials and Methods**

1. ***Selection of strong and poor shift groups***

Individuals are known to vary considerably in their ability to shift a decision criterion, with some failing to do so despite explicit instructions, practice, and confirmed task understanding (*37-41*). Since no reliable predictors of criterion shifting exist prior to task administration (*37, 39, 41*), participants were classified post hoc into strong or poor shift groups, with poor shifters defined as ∆*c_a_* < 0.30, consistent with prior classifications of adequate criterion shifting (*27, 32*).

**Behavioral Analysis**

The recognition memory task comprised four test conditions defined by two orthogonal manipulations: decision criterion (lax versus strict) and memory strength (strong versus weak). Stimuli were drawn from four image categories (animal, food, person, plant), distributed equally across conditions and item types (old versus new). Memorability scores assigned to each image ranged from 0.60 to 0.92 and were likewise balanced across conditions, image categories, and item types. The task consisted of four study-test cycles, each comprising one study block followed by four test blocks (one per condition combination). On each trial, participants indicated their confidence as high or low. Based on the degree of criterion shift, participants were divided into two groups: good shifters (*N* = 17 sessions) and poor shifters (*N* = 12 sessions).

1. ***Accuracy***

Overall recognition accuracy did not differ significantly between old (*M* = 75.7 ± 15.3%) and new items (*M* = 73.4 ± 11.8%; *p* = .513; Fig. S1A), nor between the strong and poor shift groups (*p* = .309; Fig. S1B). In the strict condition, accuracy was significantly lower for old items (*M* = 68.9 ± 21.9%) than new items (*M* = 83.8 ± 13.0%; *p* = .002), but this pattern reversed in the lax condition, where accuracy was higher for old (*M* = 82.6 ± 15.6%) than new items (*M* = 62.9 ± 21.2%; *p* < .001), consistent with choices favoring “new” and “old” judgments, respectively (Fig. S1C, left). Under strong memory strength conditions, recognition accuracy was significantly higher for old than new items (*M* = 83.5 ± 14.8% vs. 73.2 ± 12.3%; *p* = .006), whereas this difference was absent under weak memory strength conditions (*M* = 68.0 ± 17.7% vs. 73.6 ± 12.9%; *p* = .117), demonstrating that the manipulation selectively affected recognition of old items (Fig. S1C, right).

A median split of memorability scores revealed higher recognition accuracy for high versus low memorability images (77.5% ± 10.1% vs. 71.7% ± 9.4%; *p* < .001), with no significant difference between old and new items across memorability score quartiles (Fig. S1D). Accuracy did not differ significantly across test blocks or image categories in any of the four test conditions (Fig. S1, E and F). High confidence responses were significantly more accurate than low confidence responses (*M* = 82.2 ± 12.2% vs. 65.8 ± 10.5%; *p* < .001; Fig. S1G), and this pattern held across shift group, memorability scores, and test conditions (Figs. S1H-J).

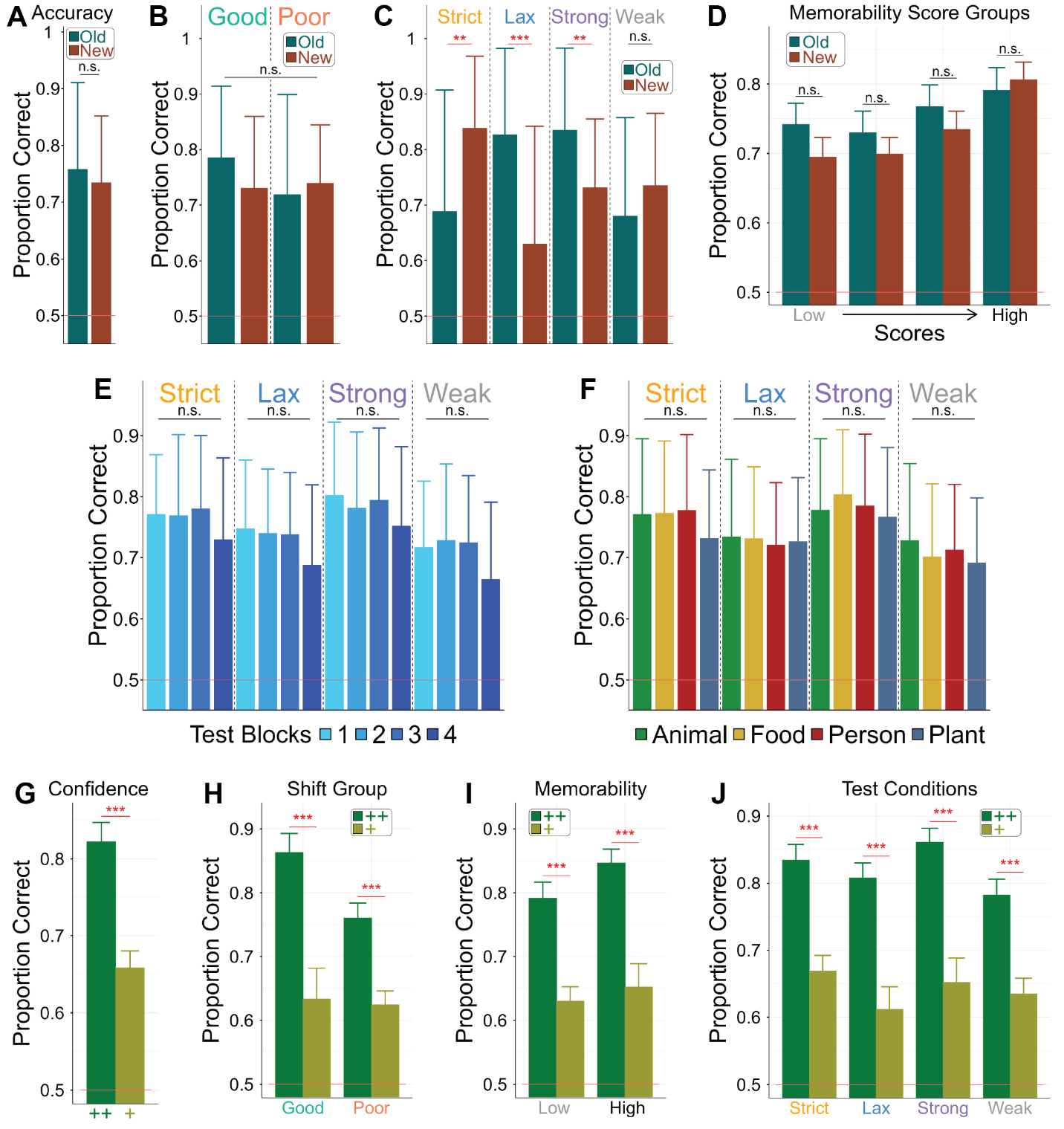

**Fig. S1. Behavioral accuracy.** (**A** to **D**) Mean ± SD proportion correct for old and new items overall (**A**), separated by shifter group (**B**), test condition (**C**), and memorability score quartile (**D**). Old/new differences in the strict and lax conditions reflect choices favoring “new” and “old” choices, respectively, while the strong memory condition shows a recognition advantage for old over new items, consistent with a benefit of repeated study. (**E** and **F**) Mean ± SD proportion correct did not differ significantly across test blocks (**E**) or image categories (**F**) within any test condition. (**G** to **J**) Mean ± SD proportion correct was significantly higher for high (++, green) than low (+, yellow) confidence responses overall (**G**), and separately by shifter group (**H**), memorability score (**I**), and test condition (**J**).

2. ***Reaction time***

Reaction times were measured from stimulus onset, encompassing the 1000 ms stimulus presentation plus the time taken to respond once the response screen appeared. For reaction time analyses, trials exceeding 3 SDs above the grand mean were excluded (1.22% of total trials). For analyses involving confidence ratings, we additionally excluded the 5 sessions in which participants did not provide at least 3 confidence rating types per response.

Overall, reaction times were significantly shorter for hits than correct rejections (*M* = 1.51 ± 0.55 s vs. 1.75 ± 0.62 s; *p* = .008), but did not differ significantly between misses and false alarms (*M* = 2.39 ± 1.16 s vs. 2.42 ± 0.96 s; *p* = .834; Fig. S2A). Reaction times for correct responses did not differ significantly between good and poor shifter groups (Fig. S2B). Reaction times were significantly shorter for high than low confidence responses (*M* = 1.48 ± 0.55 s vs. 2.49 ± 1.09 s; *p* < .001; Fig S2C), and this pattern held regardless of response type (Fig. S2D). Across test conditions, reaction times for high confidence hits did not differ significantly from high confidence correct rejections, suggesting that the overall difference between these response types was driven primarily by low confidence responses (Fig. S2E). Figure S2, F to H, shows reaction times for each response type, broken down by test block, image category, and memorability score.

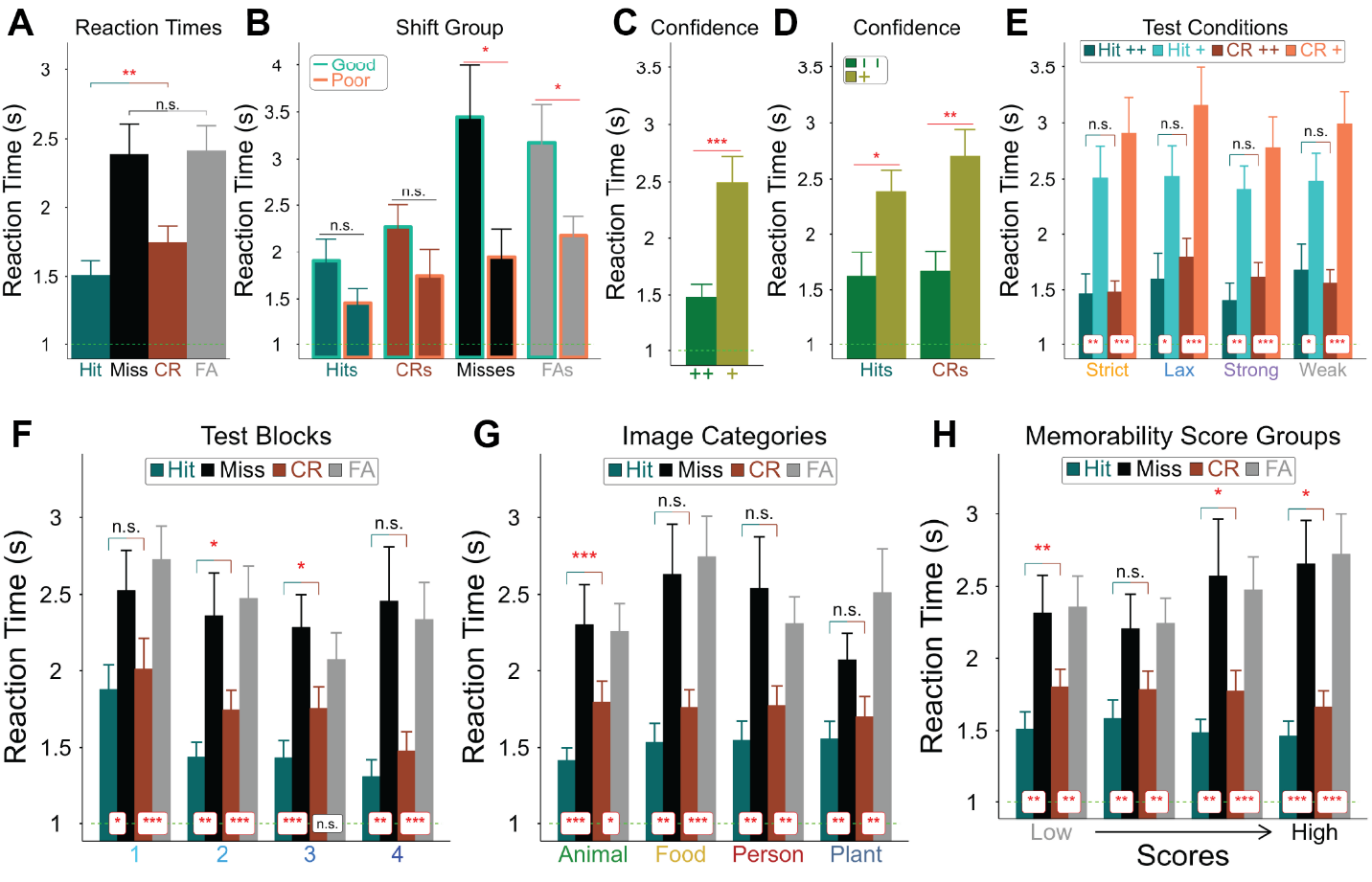

**Fig. S2. Behavioral reaction times.** A green dotted line at y = 1 s marks stimulus offset, since responses could not be made until after this point in each trial. (**A** and **B**) Mean ± SEM reaction times for hits, misses, correct rejections (CRs), and false alarms (FAs), overall (**A**) and by shifter group (**B**). (**C** and **D**) Mean ± SEM reaction times by confidence level (high [++] vs. low [+]), collapsed across response types (**C**) and separately for hits and CRs (**D**). (**E** to **H**) Mean ± SEM reaction times for each response type, broken down by test condition (**E**), test block (**F**), image category (**G**), and memorability scores (**H**).

**Neuron properties**

1. ***Firing rates during baseline and stimulus periods***

Mean firing rates during the baseline (−1000 to 0 ms before onset) and stimulus periods (200 to 1200 ms after onset) for all neurons in the amygdala, hippocampus, dACC, and pre-SMA are shown in Fig. S3A, separately for neurons with higher firing rates during the baseline versus stimulus period. On average, firing rates were higher in the MFC (4.66 ± 4.47 Hz) than in the MTL (2.67 ± 2.78 Hz; *p* < .001). This breakdown is further separated by neuron type (CS, MS, and VS) and brain region (MFC vs. MTL) in Fig. S3, B to F.

2. ***Proportion of cell types***

Of the 695 neurons analyzed in the MTL, 44.3% were VS (308/695), 12.8% were MS (89/695), and 8.3% showed a preference for the criterion condition during the baseline period (58/695), and 41.6% showed no significant modulation by any of these categories (289/695; Fig. S3, G and H). Of the 505 neurons analyzed in the MFC, 15.8% were MS (80/505), 9.9% were VS (50/505), 9.9% were CS (50/505), 13.3% showed a baseline criterion preference (67/505), and 59.4% showed no significant modulation by any of these categories (300/505; Fig. S3, I and J).

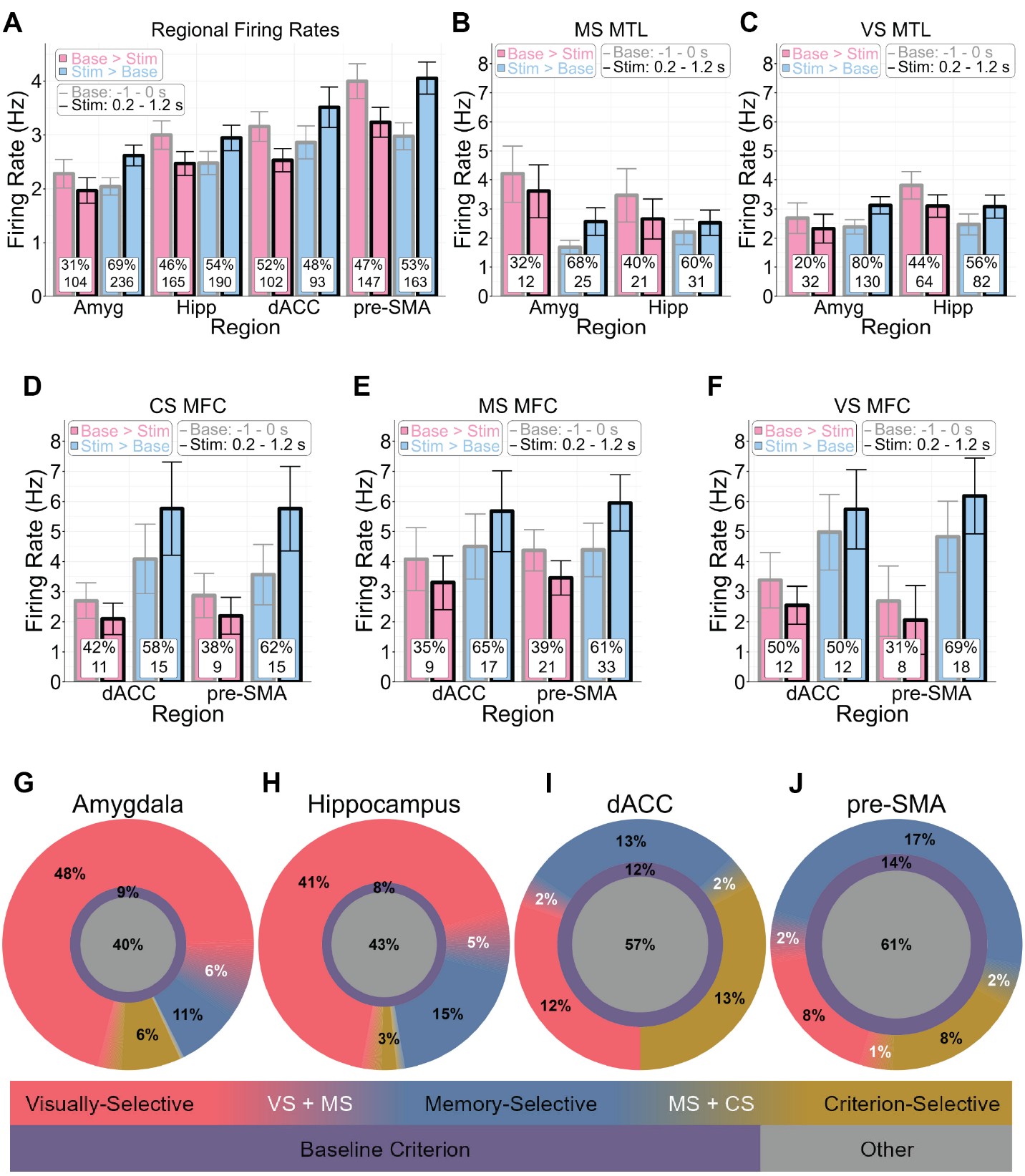

**Fig. S3. Neuronal firing rates and selective proportions across regions.** (**A**) Mean ± SEM firing rates during baseline (gray; −1000 to 0 ms before stimulus onset) and stimulus periods (black; 200 to 1200 ms after stimulus onset) for neurons with greater firing rates during the baseline (pink; Base > Stim) or stimulus period (blue; Stim > Base), shown across all neurons within the amygdala (Amyg), hippocampus (Hipp), dACC, and pre-SMA. The percentage and total count of neurons within each region showing higher firing rates during the baseline than the stimulus period is indicated at the base of each bar pair. (**B** to **F**) Mean ± SEM firing rates during baseline and stimulus periods, shown separately for MS MTL (**B**), VS MTL (**C**), CS MFC (**D**), MS MFC (**E**), and VS MFC (**F**) neurons. (**G** to **J**) Pie charts depicting the relative proportions of VS (pink), MS (blue), CS (yellow), baseline criterion condition (purple), and unclassified (gray) neurons in the amygdala (**G**), hippocampus (**H**), dACC (**I**), and pre-SMA (**J**). Percentages at color boundaries (white) indicate the proportion of neurons selective for both adjacent cell types.

**MFC criterion-selective neurons**

Time-resolved single-trial population decoding of MFC CS neurons revealed two peaks, an “early” peak preceding and a “late” peak following the peak MS neuron decoding accuracy (Fig. S4A). This temporal bracketing suggests that CS neurons encode favored versus unfavored choices most prominently both before memory evidence becomes available and after the memory signal has peaked. Consistent with this population-level observation, individual CS neurons often exhibited either the early component only (Fig. S4, B and C), the late component only (Fig. S4, D to F), or both components (Fig. S4, G to I), supporting the notion that CS activity represents a decision criterion signal that flanks the MS memory signal. Whether this bimodal temporal profile is a consistent property of CS neurons across participants and task contexts remains to be established. It is possible, however, that CS neurons could be further subdivided based on the timing of their differential firing between favored and unfavored choices.

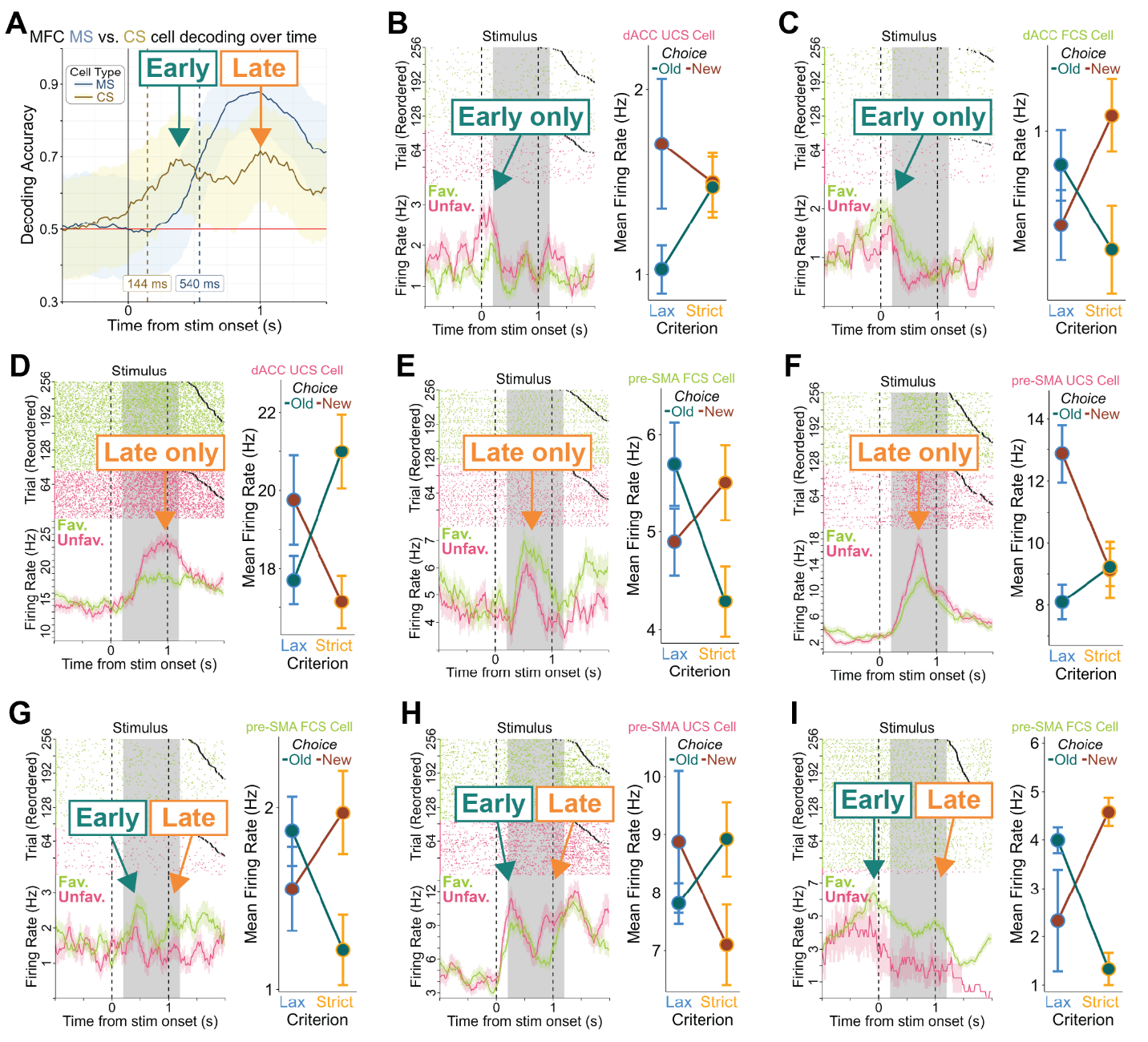

**Fig. S4. Criterion-selective (CS) neurons.** (**A**) Time-resolved single-trial decoding accuracy for MFC CS neurons (favored versus unfavored choices) and MS neurons (hits versus CRs), showing early and late CS decoding peaks flanking the MS decoding peak. (**B** to **I**) Representative MFC CS neurons are shown exhibiting the early peak only (dACC UCS, **B**; dACC FCS, **C**), the late peak only (dACC UCS, **D**; pre-SMA FCS, **E**; pre-SMA UCS, **F**), and both peaks (pre-SMA FCS, **G** and **I**; pre-SMA UCS, **H**). Raster plots (top) and peristimulus time histograms (PSTHs, bottom) show favored (lime green) and unfavored (pink) choices (left). Trials are sorted by choice type and then by reaction time (fastest to slowest); black crosses (×) mark response time. The stimulus period used for analysis (200 to 1200 ms after stimulus onset) is highlighted in each raster and PSTH. Mean ± SEM firing rates during the stimulus period show a choice-by-criterion interaction between “old” (green) and “new” (red) choices under the lax (blue) and strict (orange) criterion conditions (right).

3. ***Memory-selective and visually-selective neurons***

Memory-selective neurons in both the MTL and MFC tended to scale with memory strength, as indexed by confidence ratings and accuracy, with FS neurons firing most strongly for high confidence hits, followed by low confidence hits, errors, low confidence correct rejections, and high confidence correct rejections, whereas NS neurons showed the reverse pattern (Fig. S5, A to I), suggesting that MS neurons in both regions share similar response properties. Visually selective neurons were substantially more prevalent in the MTL than the MFC and were unaffected by either the decision criterion or memory strength (Fig. S6, A to L).

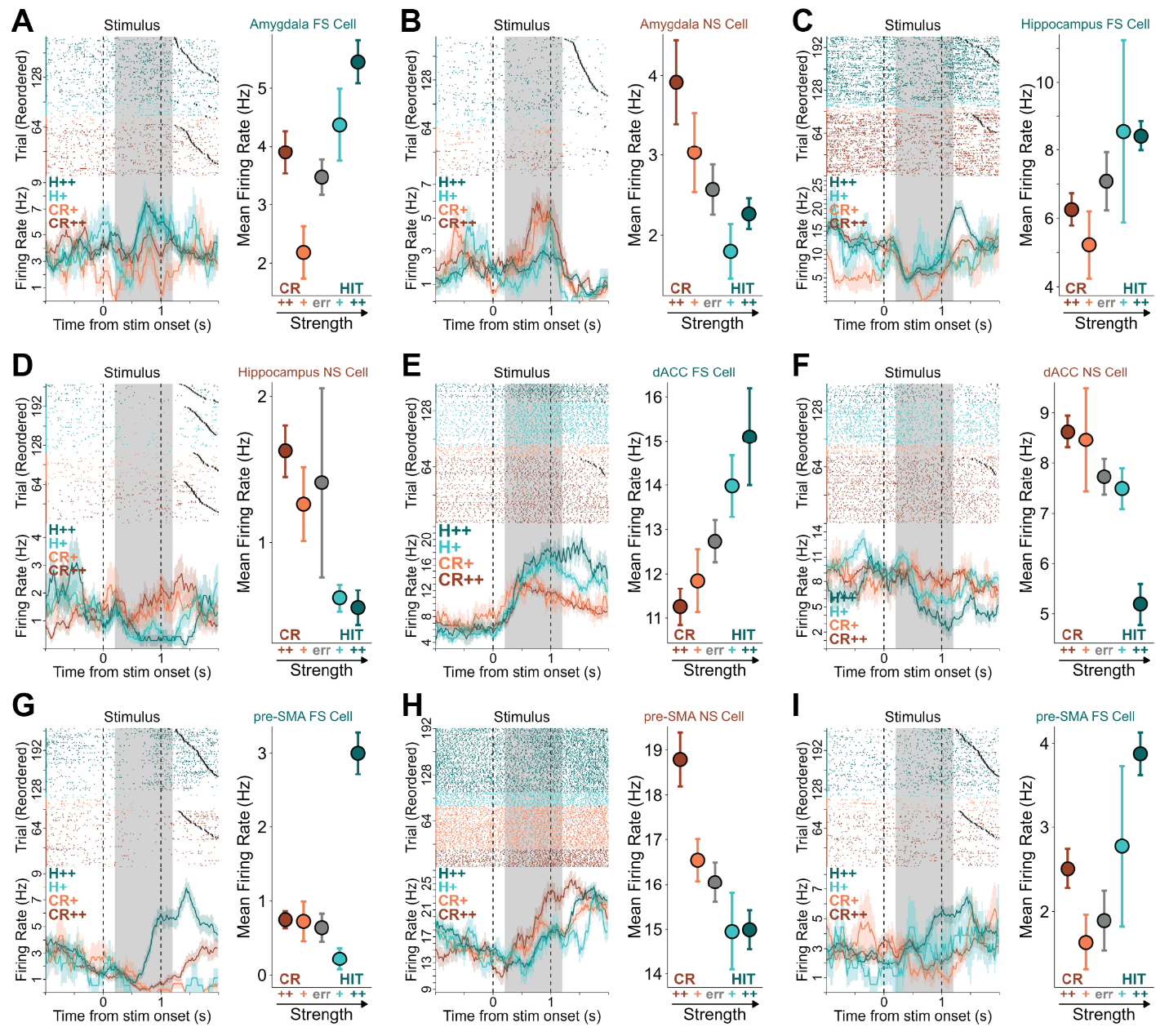

**Fig. S5. Memory-selective (MS) neurons.** (**A** to **I**) Representative MS neurons from the amygdala (FS, **A**; NS, **B**), hippocampus (FS, **C**; NS, **D**), dACC (FS, **E**; NS, **F**), and pre-SMA (FS, **G** and **I**; NS, **H**). Raster plots (top) and peristimulus time histograms (PSTHs, bottom) show high confidence hits (H++, green), low confidence hits (H+, cyan), low confidence correct rejections (CR+, coral), and high confidence correct rejections (CR++, red) (left). Trials are sorted by choice type and then by reaction time (fastest to slowest); black crosses (×) mark response time. The stimulus period used for analysis (200 to 1200 ms after stimulus onset) is highlighted in each raster and PSTH. Mean ± SEM firing rates during the stimulus period show a general monotonic change across memory strength, from high confidence hits to low confidence hits to errors (gray) to low confidence correct rejections to high confidence correct rejections, or the reverse for NS neurons (right).

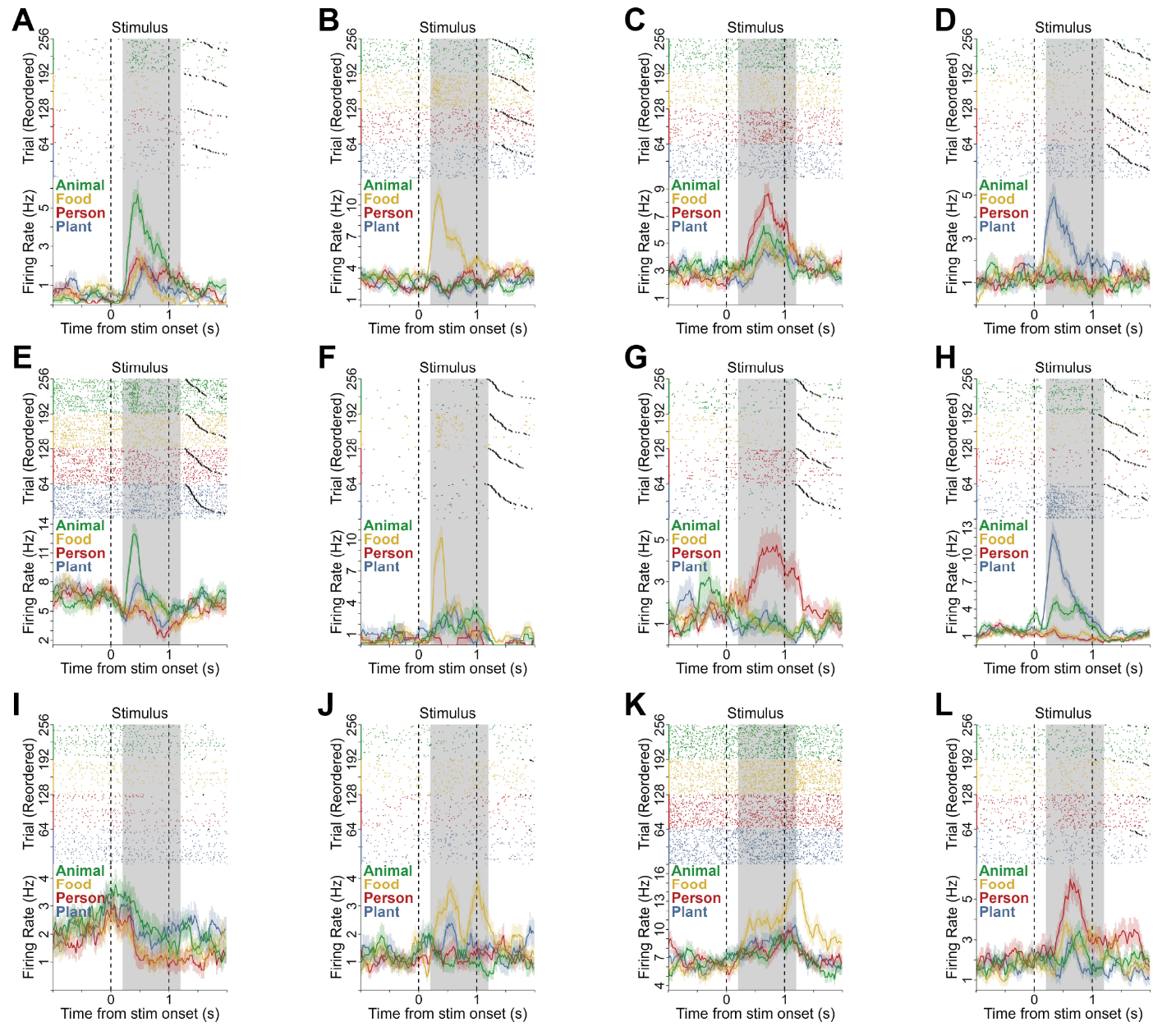

**Fig. S6. Visually-selective (VS) neurons.** (**A** to **L**) Representative VS neurons from the amygdala (**A** to **D**), hippocampus (**E** to **H**), pre-SMA (**I** and **J**), and dACC (**K** and **L**). Raster plots (top) and peristimulus time histograms (PSTHs, bottom) show trials in which the image category was of an animal (green), food (yellow), a person (red), or a plant (blue). Trials are sorted by image category and then by reaction time (fastest to slowest); black crosses (×) mark response time. The stimulus period used for analysis (200 to 1200 ms after stimulus onset) is highlighted in each raster and PSTH.

| **Patient ID** | **Age** | **Sex** | **#Ses** | **Epilepsy Diagnosis** | **Shift Group** | **∆*c_a_*** | **Strong *d_a_*** | **Weak *d_a_*** | **#Amyg** | **#Hipp** | **#dACC** | **#pre-SMA** |
| --- | --- | --- | --- | --- | --- | --- | --- | --- | --- | --- | --- | --- |
| P76CS | 24 | F | 1 | not localized | good | 1.11 | 3.06 | 2.39 | 3 | 6 | 11 | 4 |
| P78CS | 54 | F | 1 | R temporal | poor | 0.15 | 0.77 | 0.16 | 16 | 11 | 0 | 0 |
| P79CS | 42 | F | 1 | R temporal | good | 1.14 | 2.96 | 2.33 | 21 | 12 | 13 | 7 |
| P80CS | 24 | F | 1 | not localized | poor | -0.04 | 2.34 | 1.43 | 28 | 7 | 11 | 18 |
| P81CS | 28 | F | 1 | L temporal | poor | 0.06 | 1.76 | 0.68 | 6 | 6 | 3 | 13 |
| P82CS | 42 | M | 1 | Bitemporal | poor | 0.03 | 2.65 | 1.49 | 17 | 23 | 9 | 1 |
| P86CS | 39 | F | 2 | not localized | good | 3.64 | 1.96 | 0.81 | 20 | 6 | 34 | 31 |
| P92CS | 30 | F | 1 | L occipital | good | 0.61 | 1.22 | 0.93 | 13 | 10 | 6 | 32 |
| P97CS | 32 | M | 1 | R temporal | good | 0.34 | 1.83 | 1.62 | 28 | 20 | 11 | 5 |
| P98CS | 30 | F | 1 | Bitemporal | poor | 0.09 | 1.30 | 0.84 | 21 | 25 | 9 | 29 |
| P100CS | 25 | M | 1 | R temporal | good | 0.39 | 3.66 | 2.04 | 29 | 12 | 0 | 20 |
| P101CS | 34 | M | 1 | R temporal | good | 1.55 | 2.60 | 1.84 | 16 | 21 | 0 | 12 |
| P102CS | 55 | F | 1 | R temporal | good | 0.72 | 2.81 | 1.51 | 28 | 24 | 15 | 17 |
| P103CS | 51 | M | 1 | Bitemporal | good | 0.69 | 1.45 | 0.90 | 18 | 16 | 0 | 23 |
| P106CS | 37 | M | 3 | L temporal L frontal | good | 0.49 | 2.54 | 1.78 | 36 | 35 | 17 | 45 |
| P107CS | 65 | F | 1 | L temporal | good | 0.86 | 0.35 | 0.62 | 20 | 10 | 37 | 45 |
| TWH219 | 28 | M | 1 | L frontal | poor | 0.00 | 1.31 | -0.15 | 0 | 2 | 4 | 12 |
| TWH222 | 43 | M | 3 | R temporal | poor | 0.05 | 1.50 | 0.55 | 17 | 2 | 0 | 0 |
| TWH227 | 19 | M | 3 | R temporal R occipital | good | 0.47 | 1.09 | 0.52 | 3 | 38 | 2 | 13 |
| TWH228 | 34 | F | 3 | L temporal L occipital | poor | 0.05 | 2.00 | 1.39 | 0 | 69 | 0 | 0 |

**Table S1**. Patient demographics, pathology, behavioral metrics, and neuron counts (aggregated across sessions where applicable). Abbreviations: #Ses, number of sessions conducted; R, right hemisphere; L, left hemisphere; #Amyg, #Hipp, #dACC, and #pre-SMA, number of neurons recorded in the amygdala, hippocampus, dorsal anterior cingulate cortex, and pre-supplementary motor area, respectively.
